# A multi-functional small molecule alleviates fracture pain and promotes bone healing

**DOI:** 10.1101/2022.05.27.493755

**Authors:** Yu-Ru V. Shih, David Kingsley, Hunter Newman, Jiaul Hoque, Ankita Gupta, B. Duncan X. Lascelles, Shyni Varghese

**Affiliations:** Department of Orthopaedic Surgery, Duke University School of Medicine; Durham, NC 27710, USA; Department of Mechanical Engineering and Materials Science, Duke University; Durham, NC 27710, USA; Translational Research in Pain Program, Department of Clinical Sciences, College of Veterinary Medicine, North Carolina State University, Raleigh, NC 27607, USA; Thurston Arthritis Center, University of North Carolina School of Medicine, Chapel Hill, NC 27599, USA; Center for Translational Pain Medicine, Department of Anesthesiology, Duke University School of Medicine, Durham, NC 27710, USA; Comparative Pain Research and Education Center, College of Veterinary Medicine, North Carolina State University, Raleigh, NC 27607, USA; Department of Biomedical Engineering, Duke University; Durham, NC 27710, USA

## Abstract

Skeletal injuries are a major cause of morbidities worldwide with bone fractures accounting for a substantial portion. Patients suffering from bone fractures and undergoing surgery experience different levels of pain throughout the healing process requiring pain-mitigating interventions. Furthermore, a considerable number of bone fractures suffer from delayed healing, and unresolved acute pain may transition to chronic and maladaptive pain. Current management of pain involves treatment with NSAIDs and opioids, however, these analgesics have substantial drawbacks including delaying healing, systemic side effects, and potential for addiction. Hence, a therapeutic approach that concomitantly attenuates pain locally and actively promotes healing would address a significant clinical problem and improve the overall functional outcome for patients. Herein, we tested the hypothesis that the purine molecule, adenosine, could simultaneously alleviate fracture pain and promote healing by targeting different adenosine receptor subtypes in different cell populations. Our results demonstrate that local delivery of adenosine inhibited nociceptive activity of peripheral neurons through activation of adenosine A1 receptor (ADORA1) and mitigates pain. Concurrently, localization of adenosine at the fracture site also promoted osteogenic differentiation of mesenchymal stromal cells through adenosine A2B receptor (ADORA2B) and improved bone healing. Although further work is needed to extend the findings to human patients, this study provides evidence that the unique functional properties of adenosine along with its local delivery could provide an innovative, safe, and translatable therapeutic strategy to treat bone trauma and associated pain.

**One Sentence Summary:** Adenosine as a therapeutic for fracture pain and healing

## INTRODUCTION

Bone fracture is a common injury associated with sports, accidents, or falls. These injuries and post-surgical intervention to repair bone are accompanied by acute pain (*1*), which gradually diminishes as bone heals (*2*). The management of bone injuries generally involve stabilization, surgery, rehabilitation, administration of analgesics such as nonsteroidal anti-inflammatory drugs (NSAIDs) or opioids and allowing the injured tissue to heal (*1, 3, 4*). Despite the regenerative capacity of bone, some fractures suffer delayed healing or nonunion, typically requiring repeated surgical interventions, which is a challenging clinical problem and leads to long-term morbidity and chronic pain (*5-10*). The incidence of nonunion varies from 1.9% to 30% depending upon the severity of fracture, comorbidities, and lifestyle habits (*5-10*). In addition, the ongoing pain associated with impaired healing often transition to chronic and maladaptive pain (*11, 12*).

During bone healing, peripheral nerves play an important role; extensive sensory and sympathetic nerve sprouting occur after injury/surgery (*13*) and their inhibition negatively affects fracture healing (*14*). Paradoxically, the growth and activity of peripheral nerves also contribute to pain (*11, 13*). These changes in peripheral nerves could be attributed to an increase in neurotrophins, neuropeptides, and inflammatory molecules that play key roles in fracture healing but also contribute to induction, sensitization, and maintenance of pain (*15-17*). As a result, bone trauma is associated with significant pain requiring use of analgesics (*2, 18, 19*). While analgesics such as NSAIDs and opioids are effective to manage acute pain, their use often leads to unwarranted side effects. In addition to interfering with healing, use of NSAIDs and opioids can also result in gastrointestinal irritation and constipation, respectively (*9, 20-22*). Most importantly, the post-operative use of opioids can result in dependance and has been a driving factor behind the current opioid epidemic (*9, 23, 24*). In light of these problems with current analgesics and lack of active treatments to promote bone repair, new approaches that simultaneously manage pain *and* promote healing would address a significant clinical need. To this end, we examine a new therapeutic solution— localization of adenosine— and its efficacy to mitigate both pain and promote tissue regeneration following bone injury or orthopedic surgery.

Adenosine is a naturally occurring nucleoside, and extracellular adenosine regulates tissue function by activating the G-protein-coupled adenosine receptors - A1 receptor (ADORA1), A2A receptor (ADORA2A), A2B receptor (ADORA2B), and A3 receptor (ADORA3) (*25-27*). We and others have demonstrated the key role played by extracellular adenosine in bone health and regeneration (*28-30*), and studies have demonstrated impaired fracture healing in *Cd73* (an enzyme that generates extracellular adenosine), *Adora2a*, and *Adora2b* knockout mice (*31-33*). We have recently shown that localization of extracellular adenosine at the fracture site promoted vascularization and improved fracture repair in a murine tibial fracture model (*30*). In addition to its role in bone repair, adenosine also exhibits analgesic effects and perturbation of adenosine signaling has been shown to play a role in inflammatory and neuropathic pain (*34*). Activation of ADORA1 and ADORA3 on nociceptors, and ADORA3 on spinal microglia have been shown to elicit analgesic effects (*35, 36*). Targeting pain at the origin of noxious stimuli by local delivery of analgesics could avoid side effects while providing pain relief (*37-39*). In this study, we demonstrate that ADORA1 is involved in functional activity of dorsal root ganglion neurons and ADORA2B promotes osteogenic differentiation of mesenchymal stromal cells (MSCs), and biomaterial-assisted local delivery of adenosine to the fracture site alleviate fracture/post-surgical pain while promoting bone healing.

## RESULTS

### Peripheral neurons innervating fractured bone exhibit enhanced nociceptive phenotype and activity

Injury to bone has been shown to elicit spontaneous and palpation-induced nocifensive behaviors, reduced weight bearing, decreased rearing, and increased mechanical, thermal, and cold allodynia in mice (*19, 40, 41*). Tibial fracture-induced changes to sensory neurons were examined by harvesting the L3 and L4 DRG that innervate the tibia (*42*). Previous studies have shown that transient receptor potential cation channel subfamily V member 1 (TRPV1) expressing neurons are pivotal to nociception associated with bone trauma and cancer (*43-45*). Similarly, we found higher levels of TRPV1 expression in ipsilateral TUBB3-positive DRG neurons compared to contralateral DRG neurons by immunofluorescence staining (Fig. 1, A and B). Gene expression analyses by quantitative real-time polymerase chain reaction (RT-qPCR) also demonstrated a marked increase of *Trpv1* gene expression in ipsilateral DRG compared to contralateral DRG (Fig. 1C). Functional changes in sensory neurons were evaluated by calcium imaging. Dissociated DRG neurons from injured animals 5 days after fracture were cultured *in vitro* for 1 day and stimulated with the TRPV1 agonist capsaicin (Fig. 1D). The TRPV1 agonist capsaicin is commonly used to assess the functional responsiveness of nociceptive neurons *in vitro (46*). DRG neurons in all subsequent imaging experiments were also stimulated with capsaicin. Results showed that the average and peak intensity were significantly higher in ipsilateral DRG neurons compared to the contralateral DRG neurons (Fig. 1, E and F). These results suggest that the DRG neurons associated with the fractured limbs exhibit an elevated functional response, which could be partially associated with the increased TRPV1 expression.

**Figure 1.**
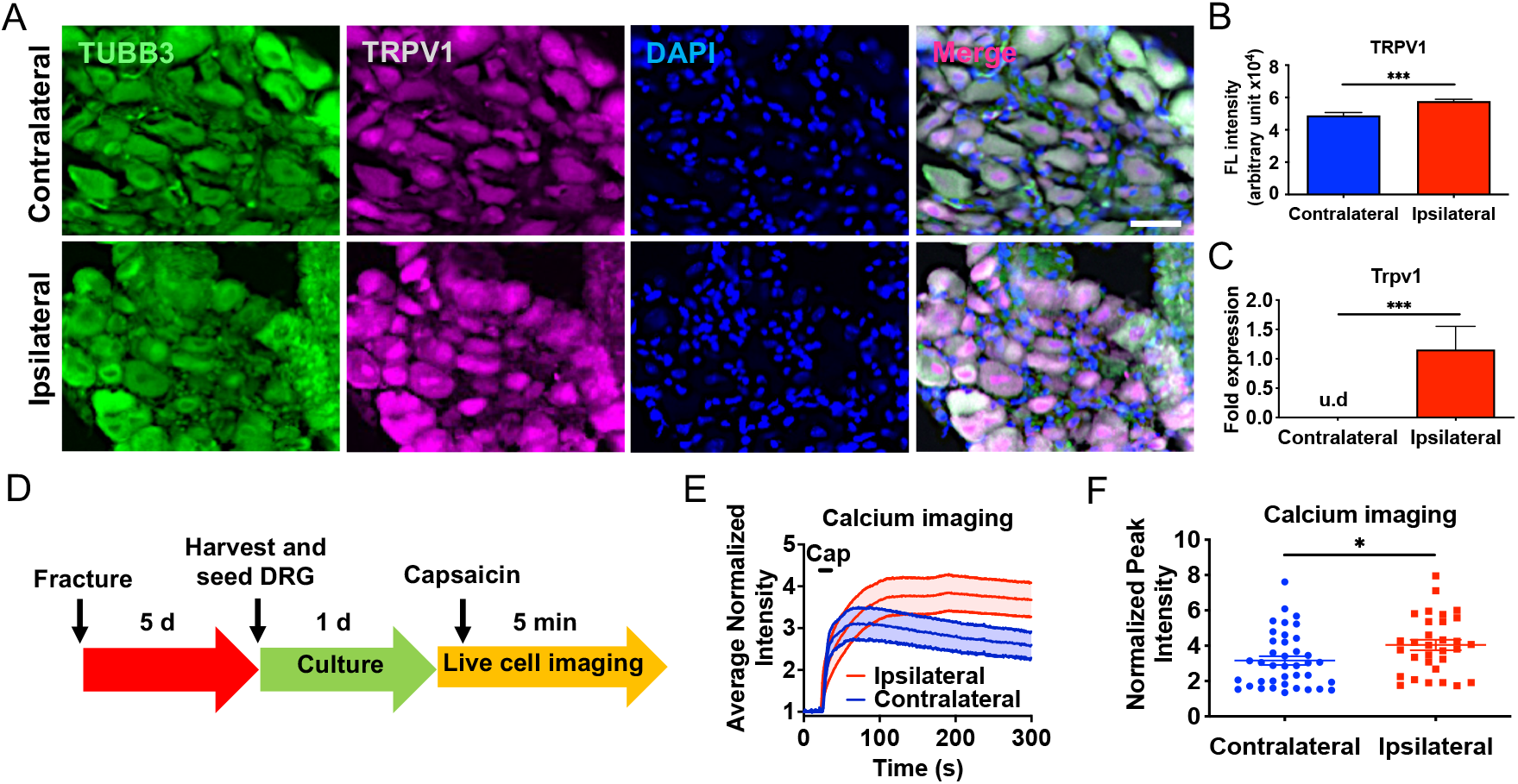
Fracture changes the phenotype and increases the functional responsiveness of DRG neurons. (**A**) Immunofluorescence staining of TUBB3 (green) and TRPV1 (violet) in ipsilateral (fractured) and contralateral (nonfractured) L3-L4 DRG from injured mice at 5 days post-fracture (dpf). DAPI stains the nucleus (blue). Scale bar, 100 μm. (**B**) Quantified TRPV1 fluorescence intensity from the immunofluorescence images (mean ± SEM, *n*>50 DRG neurons from 5 mice per group). (**C**) Relative *Trpv1* gene expression of ipsilateral and contralateral L3-L4 DRG at 5 dpf. u.d: undetectable (mean ± SEM, *n*=3 mice per group). (**D**) Experimental design of *in vitro* calcium imaging of dissociated DRG neurons from fractured mice. (**E**) Normalized average signal intensity (340/380 nm) from Fura-2 imaging of ipsilateral and contralateral dissociated DRG neurons. TRPV1 agonist capsaicin (Cap) was added at the specified time (black line) to stimulate cells and signals were normalized to baseline (*n*=31-37 cells pooled from 3 mice per group). (**F**) Normalized peak intensity from the calcium imaging of stimulated neurons (mean ± SEM). Statistical analyses were performed by two-tailed unpaired *t* test. \**P* < 0.05, \*\**P* < 0.01, \*\*\**P* < 0.001.

### Adenosine attenuates NGF-induced sensitization of DRG neurons through adenosine A1 receptor

Gene expression analyses of whole DRG showed *Adora1* is expressed at a significantly higher level than *Adora2a, Adora2b*, and *Adora3* (Fig. 2A). Immunofluorescence staining of DRG from fractured animals showed higher expression of *Adora1* in ipsilateral TUBB3-positive neurons compared to contralateral DRG neurons (Fig. 2, B and C). Gene expression levels of ipsilateral and contralateral DRG corroborated the immunofluorescence results (Fig. 2D). Fractures produce an array of inflammatory cytokines, chemokines, neurotrophins, and neurokines that have been shown to contribute to inflammatory pain (*15-17*). One such molecule is nerve growth factor (NGF) which is upregulated following fracture and its inhibition attenuates fracture pain (*14, 19*). The nociceptive effect of NGF is partially due to its role in promoting the translocation of TRPV1 to the cell surface membrane and upregulation of TRPV1 expression (*47, 48*). In line with this, we observed an upregulation of *Ngf* gene expression in whole bone marrow of fractured tibiae (fig. S1). Therefore, we used NGF as a model molecule for *in vitro* sensitization of DRG neurons and examined the effect of adenosine to attenuate the increased nociceptive phenotypic and functional changes induced by NGF. Our results demonstrate that treatment with NGF significantly increased TRPV1 receptor expression in TUBB3-positive neurons (Fig. 3, A and B). Subsequently, calcium imaging was performed on dissociated DRG neurons in the presence or absence of incubation with NGF, followed by treatment with adenosine or control (Fig. 3C). Results showed that the exposure of DRG neurons to NGF increased their functional responses to capsaicin, which was attenuated by adenosine (fig. S2, Fig. 3, D and E). Note that adenosine alone decreased the baseline functional responses of control neurons. The adenosine concentration used was identified from an initial screening (fig. S3, A and B). To determine whether the ADORA1 was responsible for adenosine-mediated decrease in functional responses, calcium imaging was performed on dissociated DRG exposed to NGF and treated with adenosine in the presence or absence of ADORA1 inhibitor 8-Cyclopentyl-1,3-dipropylxanthine (DPCPX) (Fig. 3F). Results showed an increase in the average and peak fluorescence intensity upon treatment with DPCPX compared to the vehicle control lacking DPCPX (Fig. 3, G and H). We further confirmed the effects of adenosine on neuronal sensitivity by measuring the membrane potential of dissociated DRG neurons *in vitro* using FluoVolt, a voltage-sensitive dye (fig. S4). FluoVolt imaging showed that the NGF exposure increased the membrane potential, but the increase was attenuated in the presence of adenosine (fig. S4, A to C). We also examined the effect of short term (1 h) exposure of NGF on DRG neurons by measuring calcium flux changes in the presence or absence of adenosine (fig. S5A). The results showed NGF significantly increased the average and peak calcium intensity, which was attenuated in the presence of adenosine (fig. S5, B to D).

**Figure 2.**
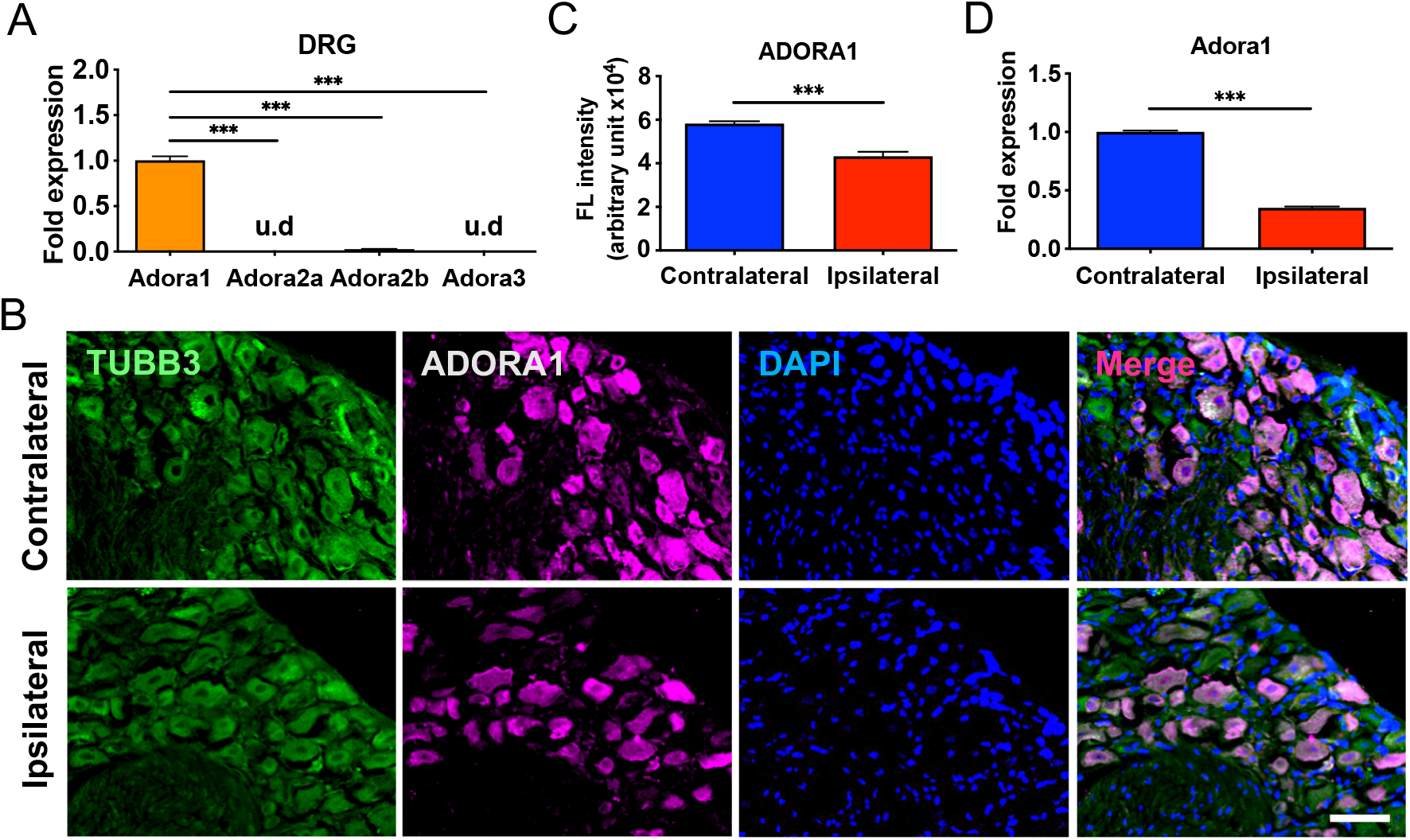
DRG neurons express adenosine A1 receptor. (**A**) Relative gene expression of adenosine receptors in L3-L4 DRG (mean ± SEM, *n*=3 mice pooled together. One-way analysis of variance [ANOVA] with Tukey post hoc test). u.d: undetectable. (**B**) Immunofluorescence images of TUBB3 (green) and ADORA1 (violet) in ipsilateral and contralateral L3-L4 DRG from fractured mice at 5 days post-fracture (dpf). DAPI stains nucleus (blue). Scale bar, 100 μm. (**C**) Quantified ADORA1 fluorescence intensity from the immunofluorescence images (mean ± SEM, *n*>50 neurons from 4 mice per group. Two-tailed unpaired *t* test). (**D**) Relative *Adora1* gene expression of ipsilateral (fractured) and contralateral (nonfractured) L3-L4 DRG at 5 dpf (mean ± SEM, *n*=7 mice pooled together per group. Two-tailed unpaired *t* test). \*\**P* < 0.01, \*\*\**P* < 0.001.

**Figure 3.**
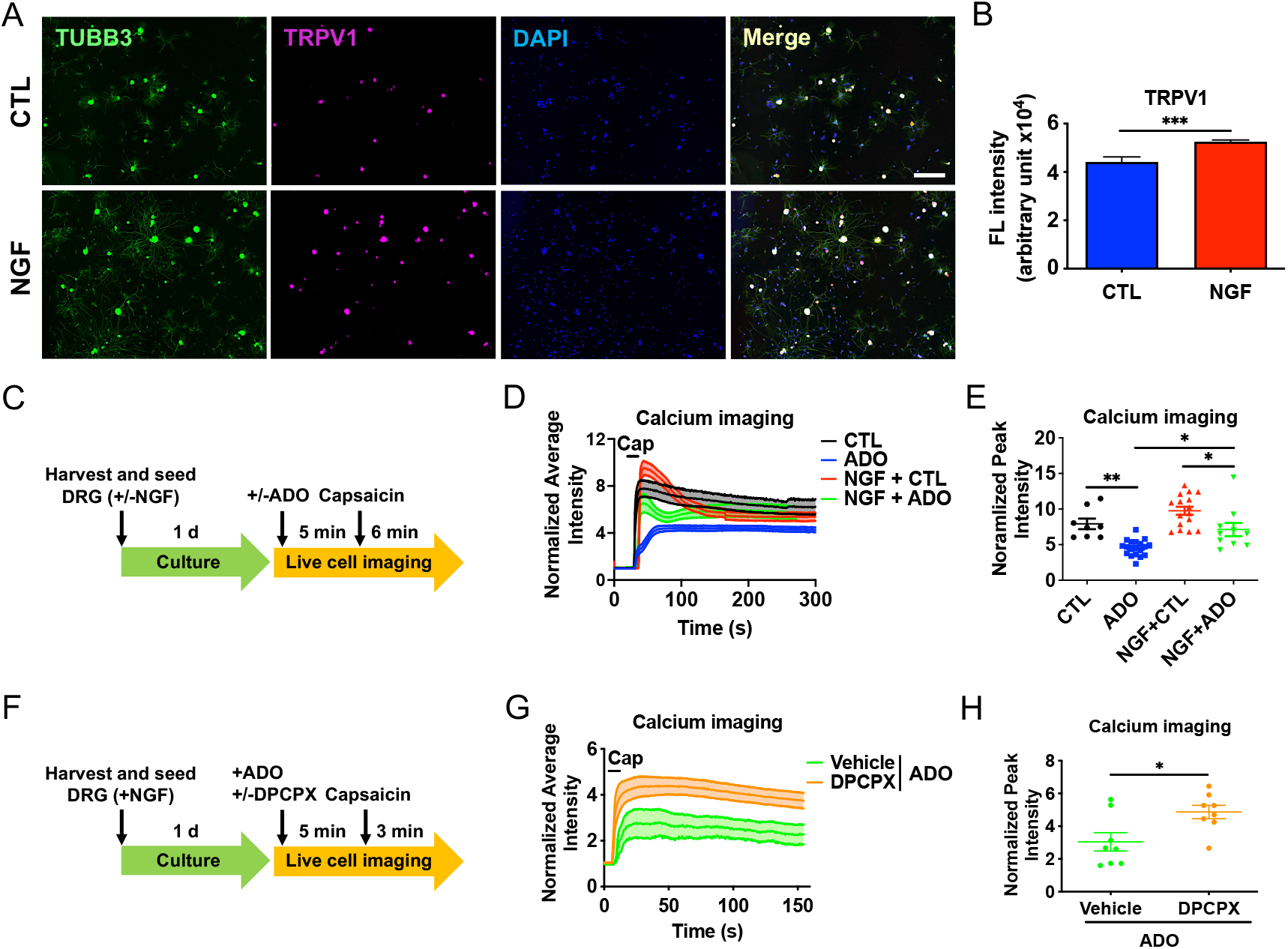
Adenosine mitigates DRG activity through adenosine A1 receptor. (**A**) Immunofluorescence staining of TUBB3 (green) and TRPV1 (violet) in dissociated DRG neurons treated with or without nerve growth factor (NGF; 200 ng/mL). CTL: control. DAPI stains nucleus (blue). Scale bar, 200 μm. (**B**) Quantified TRPV1 fluorescence intensity from the immunofluorescence images (mean ± SEM, *n*>40 neurons per group. Two-tailed unpaired *t* test). (**C**) Experimental design of calcium imaging. 1d: 1 day culture; ADO: adenosine. (**D**) Normalized average signal intensity from Fura-2 calcium imaging. TRPV1 agonist capsaicin (Cap) was added at the specified time (black line). (**E**) Normalized peak intensity from Fura-2 calcium imaging (mean ± SEM, *n*=8-16 cells per group. Two-way ANOVA with Tukey post hoc test). (**F**) Experimental design used to examine the role of ADORA1 on nociception of dissociated DRG neurons. (**G**) Normalized average intensity of DPCPX cultures compared to the vehicle control (DMSO) using calcium imaging. TRPV1 agonist capsaicin (Cap) was added at the specified time (black line). (**H**) Normalized peak intensity from the calcium imaging (mean ± SEM, *n*=8-16 cells per group. Two-tailed unpaired *t* test). \**P*<0.05, \*\**P* < 0.01, \*\*\**P* < 0.001.

### Local delivery of adenosine in fractures improves weight bearing

The *in vitro* results demonstrate the ability of adenosine to inhibit functional responses of DRG neurons including those sensitized by the fracture-associated pro-nociceptive neurokine NGF. We next examined the pain-relieving effect of adenosine in animals with fracture trauma by local delivery of adenosine following fracture. To aid local delivery, we used poly(ethylene glycol)-co-6 aminocaproic acid macroporous hydrogels functionalized with 3-(acrylamido)phenylboronic acid (3APBA), where the PBA moieties were used to load the adenosine molecules (*30*). Details about the fabrication of the macroporous hydrogels are described in Materials and Methods. Successful synthesis of the hydrogel precursors, poly(ethylene glycol) diacrylate (PEGDA) and N-acryloyl-6-aminocaproic acid (A6ACA) from poly(ethylene glycol) (PEG) and 6-aminocaproic acid (6ACA) respectively, was assessed by FTIR (fig. S6, A and B) and NMR (fig. S7, A and B). The fabricated macroporous hydrogel (PEG-6ACA-PBA) was characterized by Fourier transform infrared (FTIR) spectroscopy (fig. S8) and proton nuclear magnetic resonance (^1^HNMR) (fig. S9) spectroscopy. Adenosine loading was determined by UV/Vis spectrophotometer at a wavelength of 260 nm, which reveals that ∼0.26±0.04 mg adenosine was incorporated per 1 mg of the macroporous hydrogel. The *in vitro* release kinetics of adenosine from the macroporous hydrogels showed an initial rapid release followed by a continuous release (fig. S10). Following fracture surgery, the animals were treated with the macroporous hydrogels with or without adenosine.

Weight bearing of mice was measured to examine the effect of adenosine on mitigating fracture pain. The experimental timeline is described in Fig. 4A. Fracture surgery was performed in the absence of analgesia and the post-fracture site was treated immediately with the macroporous hydrogels measuring approximately 4 mm (length) X 2 mm (width) X 1 mm (thick) containing either 1.05±0.1 mg adenosine (ADO) or no adenosine (CTL). Hindlimb weight bearing was of the hindlimb weight placed on both hindlimbs. Tibial fracture resulted in a significant reduction in ipsilateral weight bearing percentage in the control group at 2, 4, and 8 days post-fracture compared to pre-fracture (baseline) (Fig. 4B). On the other hand, the ipsilateral weight bearing of adenosine-treated limbs was significantly decreased at 2 days post-fracture and returned back to the baseline by 8 days. When comparing weight bearing of adenosine-treated limbs to that of the untreated, fractured control animals, weight bearing of adenosine-treated measured using an incapacitance meter and ipsilateral weight bearing expressed as a percentage limbs was significantly higher compared to control-treated limbs at all post-fracture timepoints. Furthermore, the positive impact of adenosine treatment on weight bearing was associated with a large standardized effect size at each timepoint (Fig. 4C). The percent change from baseline in weight bearing was also calculated (Fig. 4D). Results showed both groups displayed significant decreases in weight bearing from the baseline at 2 and 4 days post fracture, but the degree of decreased weight bearing was less in the adenosine treated group. At 8 days post-fracture, weight bearing in the control group was still significantly reduced compared to baseline, but weight bearing in the adenosine treated group had returned to baseline levels. (Fig. 4D). To determine whether the locally delivered adenosine resulted in increased systemic blood concentrations, we measured adenosine in the peripheral blood 3 days after its implantation at the fractured tibia and found no significant increase in the adenosine level (fig. S11).

**Figure 4.**
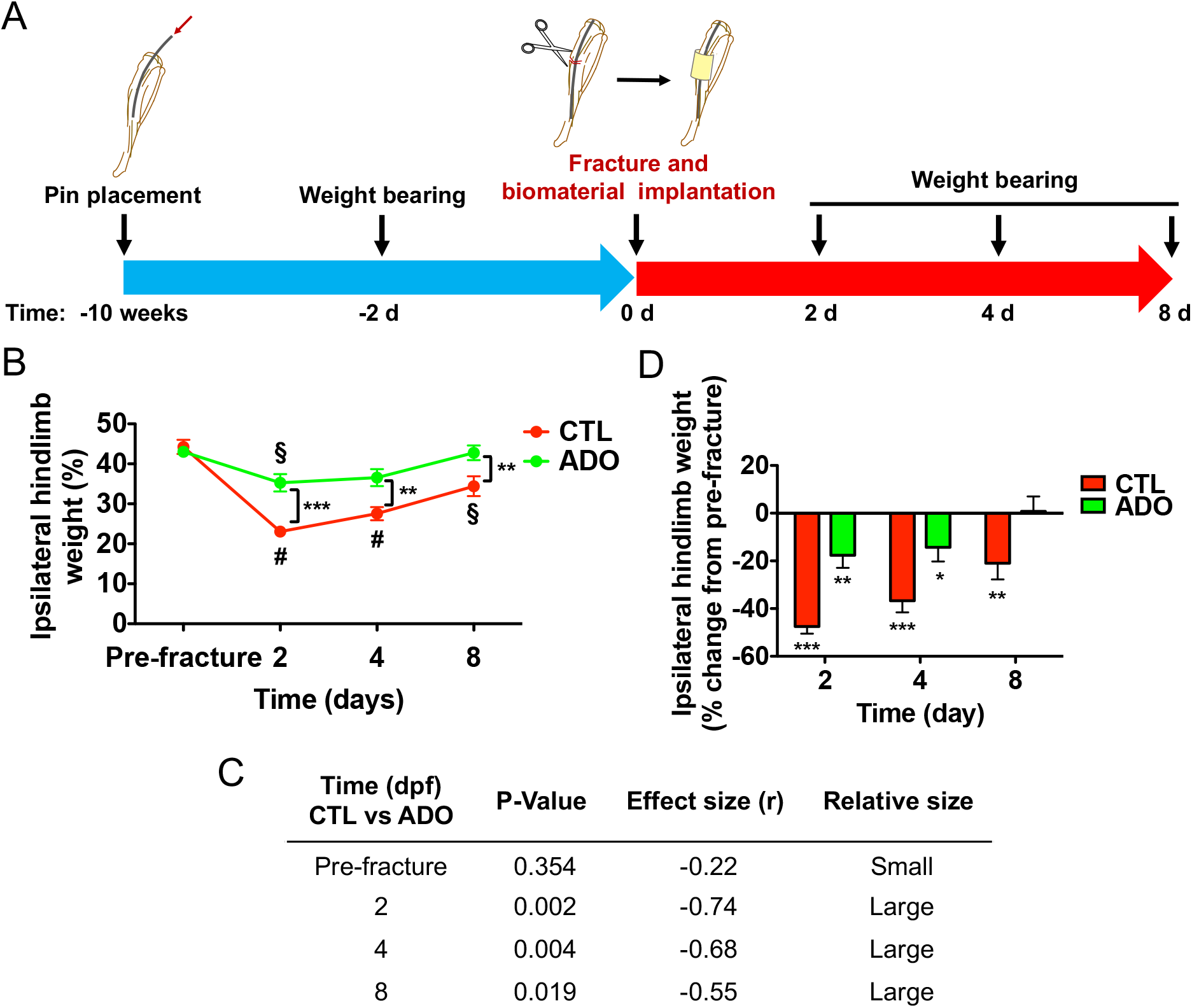
Local delivery of adenosine improves weight bearing of injured limbs. (**A**) Schematic illustrating the timeline of pin placement for stabilization, fracture surgery, biomaterial implantation, and weight bearing experiments. (**B**) Percentage of ipsilateral hindlimb weight bearing in fractured mice treated with control (CTL) or adenosine (ADO)-loaded macroporous hydrogels (*n*=9 mice per group. (mean ± SEM, *n*=9 mice per group. Friedman test with Wilcoxon signed-rank test was used to compare within the same treatment cohort over time). ^§^*P* < 0.05, ^#^*P*<0.01. Mann Whitney U test was used to compare between CTL and ADO at the same timepoint. (**C**) Effect size r of treatments on weight bearing corresponding to figure B. (**D**) Percent change of hindlimb weight bearing from pre-fracture (baseline) (mean ± SEM, *n*=9 mice per group. Mann Whitney U test was used for each treatment group compared to baseline at each timepoint). \**P*<0.05, \*\**P* < 0.01, \*\*\**P* < 0.001.

### Local delivery of adenosine improves open field movement

As vertical and ambulatory activity has been used to assess pain and functional outcomes (*49*), animals were subjected to an enclosed open field and voluntary activities were recorded for a duration of 60 min. The timeline of surgery and open field activity is shown in the schematic of the experimental design (Fig. 5A). The open field test was performed at 7 days post-fracture and representative plots of activities for each animal (5 min duration between 30^th^ and 35^th^ min) were depicted by red dots representing the vertical activity or rearing, and blue lines representing the horizontal activity or ambulation (Fig. 5B). The activities for the whole duration were collected as 5 min intervals and represented as a ratio of post-fracture to pre-fracture (fig. S12). Adenosine-treated animals had higher levels of activity than the control group animals for most of the 5-min intervals including vertical activity count, vertical movement time, ambulatory activity count, ambulatory time, total distance, and lower rest time (fig. S12, A to F). The sum of activities over the whole duration also showed significantly higher vertical activity counts, vertical movement time, ambulatory activity counts, ambulatory time, total distance, and lower rest time for the cohorts treated with adenosine compared to the control group (Fig. 5, C to H, respectively). The effect of adenosine treatment was associated with large effect sizes across all of the measured parameters (Fig. 5I).

**Figure 5.**
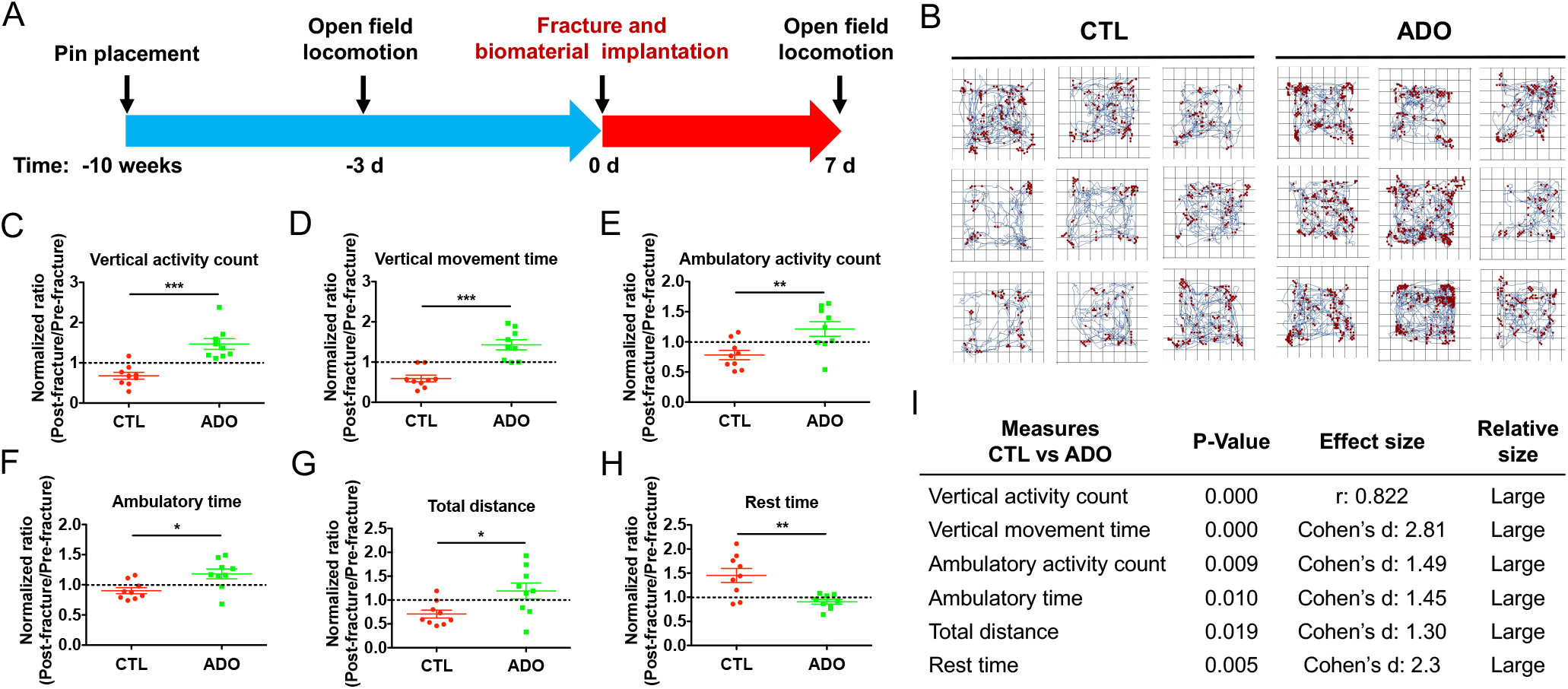
Local delivery of adenosine improves open field activity of fractured animals. (**A**) Experimental timeline of surgery, biomaterial implantation, and open field activity experiments. (**B**) Tracking of vertical activity (red dot) and horizontal activity (blue line) of fractured mice treated with control (CTL) or adenosine (ADO)-loaded macroporous hydrogels over a 5 min period at 7 dpf. Each grid represents one mouse. (**C-H**) Normalized ratio of total vertical activity count, vertical movement time, ambulatory activity count, ambulatory time, total distance, and rest time of ADO-treated mice at 7 dpf. Results are represented as a ratio of post-fracture divided by pre-fracture values (mean ± SEM, *n*=9 mice per group. Mann Whitney U test was used for vertical activity count. Two-tailed unpaired *t* test was used for other parameters). \**P*<0.05, \*\**P* < 0.01, \*\*\**P* < 0.001. (**I**) Effect size r and Cohen’s d of treatment on open field activity for each parameter in figures C-H.

### Local delivery of adenosine attenuates increased functional responsiveness of DRG neurons from mice with tibial fractures

To examine the *in vivo* effect of adenosine on the functional responsiveness of DRG neurons from mice with fractured tibiae, L3-L4 DRG were harvested 5 days post-fracture and cultured for 1 d prior to calcium imaging (Fig. 6A). Results showed that DRG neurons associated with adenosine-treated limbs had lower average and peak fluorescence intensity compared to DRG neurons from untreated, fractured control animals (Fig. 6, B and C). DRG neurons that were responsive to capsaicin had a lower but not significant percentage, in DRG from adenosine-treated animals compared to fractured controls (Fig. 6D). This finding mirrors the results from weight bearing and open field activity experiments and suggest that adenosine-mediated decrease in neuronal responsiveness is responsible for reduced pain and improved limb function in adenosine treated animals.

**Figure 6.**
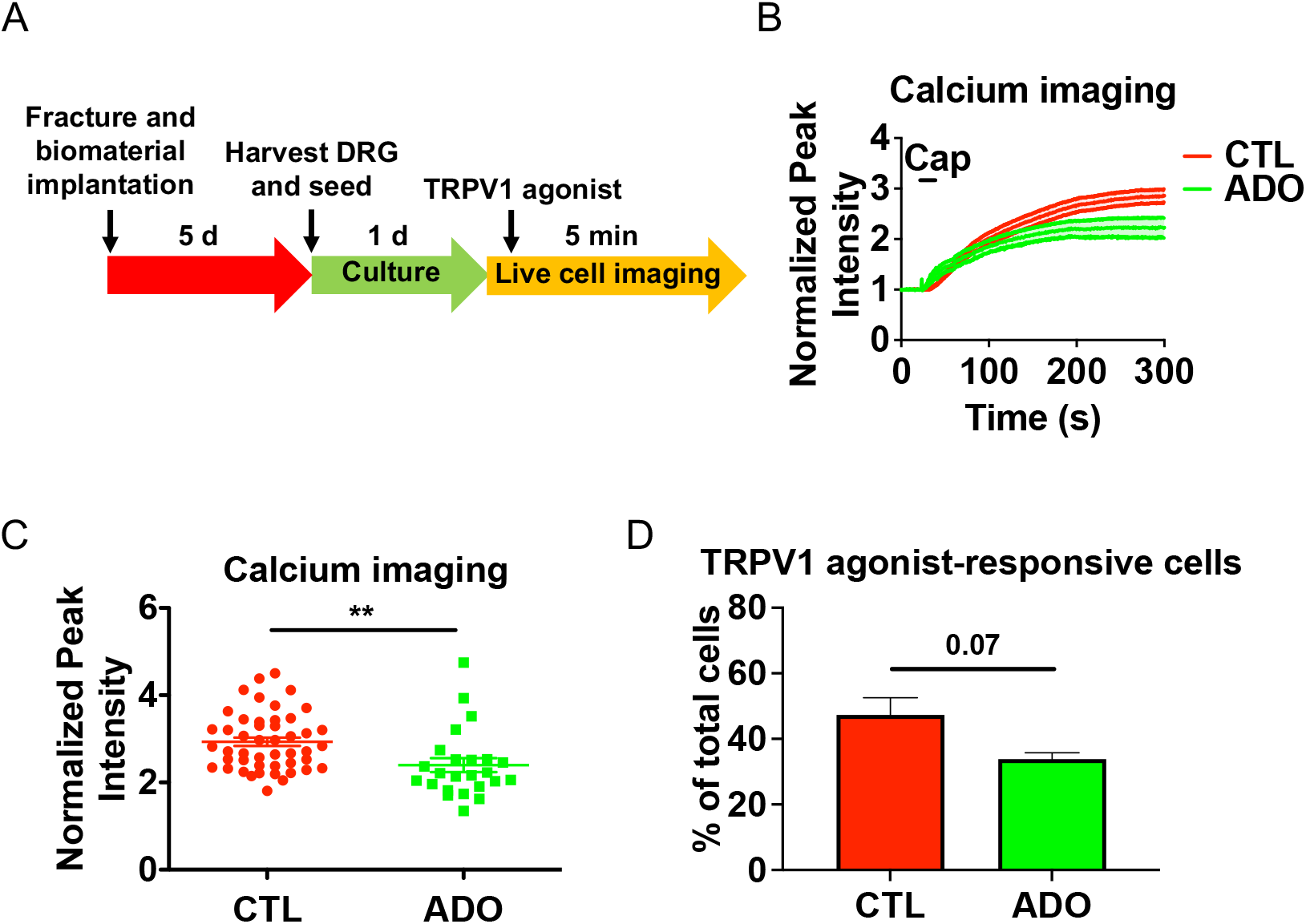
Adenosine treatment attenuates DRG activity of fractured limbs. (**A**) Experimental design for *in vitro* calcium imaging of dissociated L3-L4 DRG neurons innervating fractured limbs treated with adenosine (ADO). (**B**) Normalized average signal intensity from Fura-2 calcium imaging of DRG neurons innervating fractured hindlimbs treated with ADO and CTL groups. TRPV1 agonist capsaicin (Cap) was added at the specified time (black line) to stimulate cells and signals were normalized to baseline (*n*=24-47 cells pooled from 3 mice per group). (**C**) Normalized peak intensity from calcium imaging (mean ± SEM). (**D**) Percentage of DRG neurons from ADO and CTL group that were responsive to capsaicin (mean ± SEM). ADO: adenosine treatment. CTL: Control group without adenosine treatment. Statistical analyses were performed by two-tailed unpaired *t* test. \**P* < 0.05, \*\**P* < 0.01, \*\*\**P* < 0.001.

### Local delivery of adenosine promotes fracture healing

Local delivery of adenosine accelerated fracture healing as shown by 3D reconstructed microcomputed tomography (microCT) images (intact and cut views) and corresponding radiographs of the injured tibiae with treated groups having smaller calluses and improved healing at 21 days post-fracture, akin to our prior study (*30*) (Fig. 7A). Analyses of microCT images indicated increased bone volume and bone mineral density (Fig. 7, B and C, respectively), and lower total volume (Fig. 7D) in cohorts treated with adenosine. Similar to microCT analyses, histological images showed improved bridging across the fracture site in adenosine-treated groups compared to controls at 21 days post-fracture (Fig. 7E).

**Figure 7.**
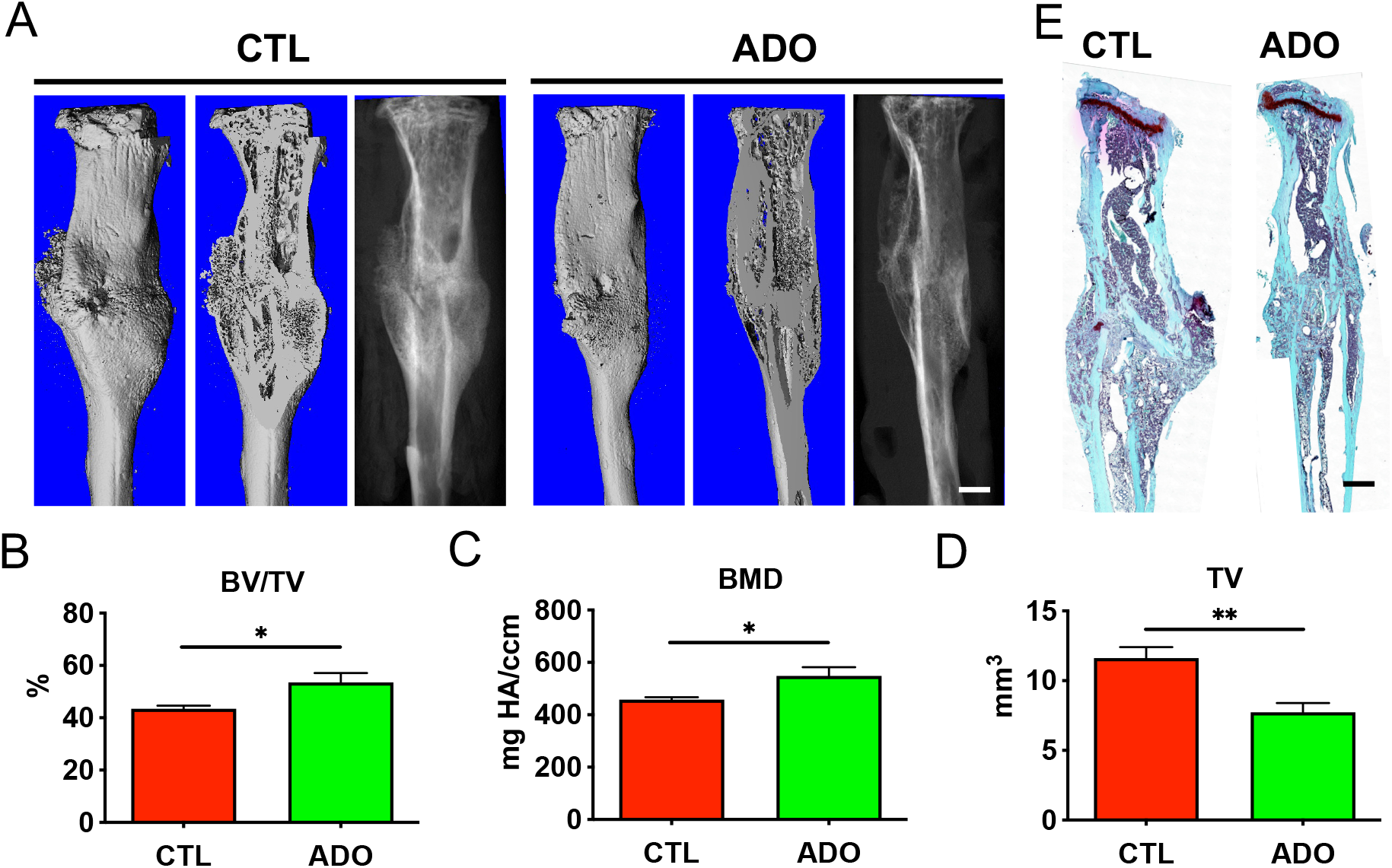
Local delivery of adenosine promotes fracture repair. (**A**) 3D reconstructed (intact and cut) and radiographs from microCT imaging of fractured tibiae treated with adenosine (ADO) or control (CTL) at 21 dpf. (**B-D**) Quantification of bone volume (BV/TV), bone mineral density (BMD), and total volume (TV) of regenerating calluses from microCT images (mean ± SEM, *n*=4 mice per group. Two-tailed unpaired *t* test). \**P* < 0.05, \*\**P* < 0.01, \*\*\**P* < 0.001. (**E**) Safranin O staining of fractured tibiae treated with adenosine (ADO) or control (CTL) at 21 dpf. Scale bar, 2 mm.

### Adenosine A2B receptor is highly expressed in osteoprogenitors of regenerating bone

Given the key role played by ADORA2B in osteogenic differentiation of stem cells (*33, 50-53*) and fracture healing (*33*), we examined the expression of ADORA2B in progenitor cells in the fracture callus. The leptin receptor (LepR) has been demonstrated to be a marker of bone marrow progenitor cells (*54*). These cells proliferate after injury and represent as a main cell source for bone tissue formation following injury (*54*). Using *Lepr-cre;tdTomato* conditional reporter mice to trace LepR(+) lineage cells, we examined the expression of ADORA2B in LepR lineage progenitors after fracture by immunofluorescent staining for ADORA2B. Our results showed abundant co-localization of ADORA2B expression in LepR lineages in the fracture calluses at 10 and 21 days post-fracture (Fig. 8A). Furthermore, gene expression analyses of adenosine receptors in bone marrow-derived MSCs demonstrated significantly higher levels of *Adora2b* expression compared to other adenosine receptor subtypes (Fig. 8B). We also examined the role of ADORA2B activation on osteogenic differentiation of MSCs. Results showed that adenosine upregulated gene expression of osteoblast transcription factors *Runx2* and *Sp7*, and osteoblast markers *Bglap* and *Ibsp* in the absence of osteogenic induction medium (Fig. 8, C-F respectively). This adenosine mediated upregulation of osteogenic markers was attenuated in the presence of ADORA2B-specific inhibitor PSB603 (Fig. 8, C-F). We also examined the expression levels of different adenosine receptors in the bone marrow and bone tissues of contralateral (unfractured) and ipsilateral (fractured) tibiae 5 days post-fracture. Results showed expression of all 4 adenosine receptors in contralateral and ipsilateral bone and bone marrow tissues and expression levels ranked from highest to lowest were: *Adora1, Adora3, Adora2b*, and *Adora2a* (fig. S13A). The expression level of *Adora2b* in the bone/marrow significantly increased during regeneration (fig. S13B).

**Figure 8.**
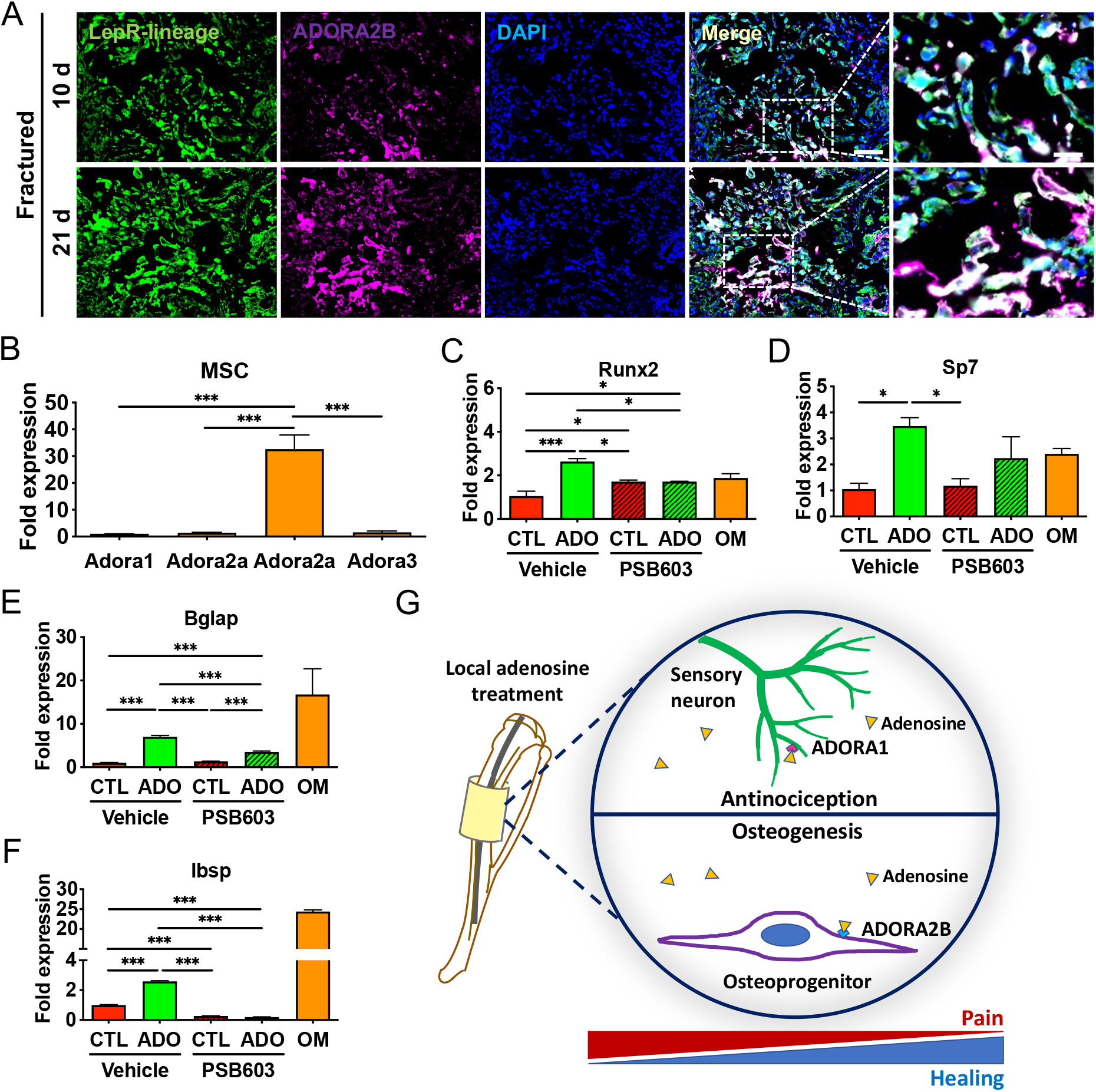
Osteoprogenitor cells express adenosine A2B receptor. (**A**) Immunofluorescence staining of Td-Tomato-labeled LepR lineage cells (green) and ADORA2B (violet) in regenerating bone of fractured mice at 10 and 21 dpf. DAPI stains nucleus (blue). dpf: days post-fracture. Scale bar, 100 μm. Magnified image scale bar, 25 μm (*n*=3 mice). (**B**) Relative gene expression of adenosine receptors in bone marrow mesenchymal stromal cells (BM-MSCs; mean ± SEM, *n*=3 mice pooled per group. One-way ANOVA with Tukey post hoc test). (**C-F**) Gene expression of Runx2, Sp7, Bglap and Ibsp in BM-MSCs, respectively. Cells were treated with adenosine (ADO, 30 µg/mL) or control (CTL) in the presence of vehicle (DMSO) or ADORA2B inhibitor PSB603 for 14 d. OM: osteogenic media. (mean ± SEM, *n*=3 pooled mice per group. Two-way ANOVA with Tukey post hoc test). \**P* < 0.05, \*\**P* < 0.01, \*\*\**P* < 0.001. (**G**) Proposed action of adenosine on multiple cell populations and adenosine receptor subtypes resulting in reduced fracture pain and improved healing.

## DISCUSSION

Fractures and associated pain (including post-surgical pain) are common clinical problems. Currently, NSAIDs and opioid analgesics are used to treat fracture pain; while these approaches are effective in managing pain, they can be associated with various side effects. Considering the limitations associated with NSAIDs and opiates, development of novel therapies to attenuate fracture pain without interfering with the tissue healing is highly desirable. In this study, we examine the potential of local delivery of adenosine to attenuate fracture pain while promoting fracture healing with improved limb function. Our findings show that adenosine can simultaneously elicit osteoanabolic and pain-mitigating functions. Specifically, adenosine activates ADORA1 on sensory neurons and inhibit neuronal excitation to mitigate pain, while stimulating ADORA2B on osteoprogenitors to promote osteogenesis and fracture healing (Fig. 8G). Although pharmacological activation of individual adenosine receptor subtypes using synthetic agonists is a viable strategy to either attenuate fracture pain or promote healing, we used adenosine due to its broader beneficial effects and ability to target multiple adenosine receptor subtypes and different cell populations.

We observed an increased TRPV1 expression in DRG neurons after injury, and higher expression and functional activity in dissociated DRG neurons sensitized by NGF, a neurokine that is elevated in the fracture microenvironment (*14, 19, 55*). TRPV1 has been shown to play a significant role in inflammatory pain (*56*), sensitizes C and Aδ fiber bone afferent neurons (*43*), and its elevated expression after bone injury correlates with elevated levels of several inflammatory molecules (*15, 16, 45*). The increased DRG neuronal function was attenuated in the presence of adenosine and pharmacological inhibition studies demonstrated the central role played by ADORA1. Studies have shown *Adora1* knockout mice experience higher sensitivity to heat and cold, and ADORA1 activation can have an analgesic effect in inflammatory and neuropathic pain (*57, 58*). This is due to ADORA1 medicated inhibition of cyclic AMP and PKA (*59, 60*), and induction of neuronal hyperpolarization by activating K^+^ channels while inhibiting Ca^2+^channels (*61-63*). Recent transcriptomic studies of human DRG neurons suggest a key role played by *ADORA1* in functional activity (*64, 65*). The established anti-nociceptive effect of ADORA1 activation along with findings that *Adora1* gene expression is significantly expressed at higher levels than other adenosine receptor subtypes in the DRG suggest it plays a dominant role in fracture pain. Our experiments revealed a decrease in *Adora1* expression in DRG and in whole bone marrow upon bone injury. A similar observation in decreased ADORA1 expression was reported in spinal cords after spinal cord injury (*66*). We speculate that the reduced expression of ADORA1 could partially be responsible for pain following injury, bestowing a protective mechanism to guard the injured tissue. However, how its expression is decreased upon injury remains elusive and warrants further investigation. Together, the relative exclusive expression of ADORA1 in DRG neurons and *in vitro* findings that adenosine inhibits NGF- and TRPV1-induced nociception through ADORA1 supports its role in adenosine-induced pain relief in fractures.

Similarly, studies have shown inflammatory molecules also directly contribute to pain, and inhibiting such molecules in itself prevents inflammatory nociception (*67, 68*). Since adenosine is a pro-resolving mediator that modulates inflammatory response of immune cells through ADORA2A, ADORA2B, and ADORA3 (*69-71*), its indirect effect on anti-nociception through regulation of inflammatory mediators cannot be ruled out. Besides inflammatory pain, adenosine has been shown to be effective against neuropathic pain (*72, 73*) and can act on both the CNS and PNS (*35, 74*), suggesting its potential for broad applications in pain relief.

Despite the potential of adenosine to mitigate pain (*75, 76*), it has not been adopted as a therapeutic due to the ubiquitous expression of adenosine receptors throughout the body, a narrow therapeutic window, and rapid clearance in circulation (*77-79*). Its short half-life and broad effects in the body suggest that adenosine was designed by nature to act locally to exert a rapid regulation of the immediate environment. With that in mind, we have leveraged a biomaterial-assisted local delivery of adenosine to regulate adenosine signaling over a prolonged period of time (*30*). Our *in vivo* studies demonstrate that local sustained delivery of adenosine attenuates fracture/surgical pain in the absence of other analgesics. Specifically, weight bearing of the injured limb treated with adenosine was significant and associated with large effect sizes. Of note, some pain from pin placement appeared to be present even after 10 weeks as ipsilateral hindlimb weight bearing percentage was 45% and not at 50% as expected, though prior studies have suggested pain returns to baseline after 4 weeks (*19*). Similarly, open field activity of animals treated with adenosine was also significantly improved and demonstrated a large effect size indicating adenosine not only reduced pain but also improved limb function. Pain diminishes after bone heals, yet the pain relief by adenosine treatment is due to its analgesic effect and not due to its effect on bone healing as we assessed the pain responses at early timepoints. Together the open field activity and weight bearing measurements at 7 and 8 days post-injury suggest sustained pain reduction and improved limb function in treated animals.

In addition to anti-nociception, the ubiquitous expression of multiple adenosine receptor subtypes present in various cell types within the bone tissue offers other potential benefits such as the ability to promote bone repair. Previous studies have shown fracture healing and osteogenic differentiation of MSCs were impaired in *Adora2b* knockout mice (*33*). However, the expression of ADORA2B in osteoprogenitors during fracture repair *in vivo* remained undetermined. In this study, we demonstrate co-localization of ADORA2B in LepR osteoprogenitors *in vivo*, further corroborating the key role of ADORA2B in MSCs to promote healing. This is consistent with our previous studies which showed adenosine promotes osteogenesis of human BM-MSCs through ADORA2B (*50, 52*). While our studies have focused on ADORA2B signaling, the contribution of ADORA2A in MSCs towards bone regeneration cannot be ruled out (*32*). As immune cells including alternatively activated (M2-like) macrophages have been shown to be important for bone repair (*80*), and A2 receptors (ADORA2A and ADORA2B) promote their polarization (*81*), it is likely that adenosine also induces a pro-regenerative immune environment by regulating immune cells through A2 receptors (*80*).

Alleviation of pain by non-addictive analgesics while not interfering with healing will have a significant clinical impact as it could reduce the use of opioids and thus prevent adverse effects of current analgesics on bone healing and potentially combating the opioid crisis. While results from the described study shows that local modulation of adenosine signaling and targeting adenosine subtypes in different cell populations in bone tissue could hold promise as an effective therapy for bone repair, additional studies need to be further explored. First, as the activity of other adenosine subtypes in inflammatory cells has been shown to reduce pain (*34*), the indirect role of adenosine receptors in immune cells to reduce pain could not be elucidated in this study. Second, the current study focused on acute pain, which clearly demonstrate a beneficial effect of adenosine, however, its effectiveness in chronic pain such as nonunions remains to be examined. Nevertheless, the unique beneficial features of adenosine to actively promote bone healing while providing pain relief offers a comprehensive and improved approach to address bone trauma and is unmatched by current therapeutics.

## MATERIALS AND METHODS

### Study design

The aim of the study was to examine the effectiveness of local delivery of adenosine in tibial bone injury to concurrently attenuate fracture/post-surgical pain while promoting healing. All animal work was performed in compliance with the National Institutes of Health (NIH) and institutional guidelines, and we followed the fracture surgery and biomaterial implantation protocols approved by the Institutional Animal Care and Use Committee at Duke University (Protocol Registry Number A151-20-07). C57BL/6J mice were used to study fracture nociception and the effect of local delivery of adenosine to mitigate pain and promote bone regeneration. *Lepr-cre; tdTomato* reporter mice were used for tracing of LepR(+) lineage cells after fracture. To determine the effect of adenosine on pain and healing, fracture surgeries were performed and treated immediately with PEGDA-6ACA-PBA macroporous hydrogels loaded with adenosine as treatment group or without adenosine as control group. Behavioral tests for pain and limb function were assessed by weight bearing and open field locomotion. A priori power analysis was used based on our pilot and previous studies to estimate the sample size requiring a statistical power (1-β) of 80% at the significance level of α = 5%. On the basis of the power calculation, the suggested number was 8 animals per group for fracture pain and 4 animals per group for fracture healing. Ten animals per group were used to study the effect of adenosine on fracture pain, and 4 animals per group were used to examine the effect of adenosine on fracture healing. The exclusion criteria for experiments are animals that died after the surgery procedures or a shift of the intramedullary pin identified after experimental endpoint post-mortem. No randomization was used and researchers were not blinded. Measurement of adenosine in the circulation was performed by using an adenosine assay. The *in vitro* functional activity of DRG neurons was assessed by calcium and membrane potential imaging and the expression of proteins were assessed by histology or immunofluorescence imaging, and gene expression by RT-qPCR. Fracture healing was analyzed by microCT and histology. The study design and timeline of animal procedures (pin placement, surgery, therapeutic intervention with and without adenosine, and behavioral tests) are illustrated in Fig. 4A (weight bearing) and Fig. 5A (open field locomotion). The sample number, statistical methods, and timepoints for each individual experiment are provided in the figure legends.

### Fracture surgery and therapeutic intervention

In all experiments, 12-to 16-week-old C57BL/6J male mice (Jackson Laboratory, Bar Harbor, ME) and *Lepr-cre; tdTomato* reporter mice were used. The average mouse weight was 25 g and mice were housed in groups of four to five, and kept on a 12-h light–dark cycle with lights off at 1800 h. All animals had *ad libitum* access to chow (PMI LabDiet 5001) and water. Fracture surgeries were performed as previously described (*30*). For procedures without hydrogel implantation, animals were anesthetized by isoflurane and injected with 1 mg/kg buprenorphine SR-LAB (ZooPharm, Laramie, WY). Each mouse was then placed in a supine position with the right tibia disinfected. After skin incision and reflection, proximal to the right knee, a 0.7-mm pin was inserted from the tibial plateau through the medullary cavity to stabilize the tibia, and a cut was made at the tibial midshaft to induce a transverse fracture. Bupivacaine (0.5%; Hospira, Lake Forest, IL) at ∼1 mg/kg body weight was applied at the surgical site following wound closure using 2 reflex clips (5 mm). For animals treated with control and adenosine-loaded macroporous hydrogels, fractures were performed as mentioned earlier but without analgesia. Hydrogels measuring approximately 5 mm (length) X 3 mm (width) X 1 mm (thick) containing either 1.05±0.1 mg adenosine (ADO) or no adenosine (CTL) were implanted adjacent to the fractures immediately. For behavioral tests, fracture surgery was delayed until 10 weeks after the pin placement to avoid interference from pain originating from sites other than the fracture site, as the pin insertion alone has been shown to induce pain (*19*). The animals were treated with the macroporous hydrogels with or without adenosine immediately following the fracture. In the cohorts for weight bearing and open field activity measurements, 1 animal from the control group and 1 animal from the treatment group were euthanized due to shift of the inserted pin following fracture.

### Statistical analyses

Statistical analyses were carried out using GraphPad Prism 9 or SPSS v.20. Normality is determined by Shapiro-Wilk test. Mann-Whitney U test and Student’s t-test were used to compare between two groups, as appropriate. Statistical analysis comparing multiple groups was performed by one-way analysis of variance (ANOVA) or two-way ANOVA with Tukey post hoc test, or Friedman test with Wilcoxon signed-rank test, as appropriate. For all comparisons, two-tailed tests were used and *P* <0.05 was considered statistically significant. Cohen’s *d* or *r* were used to calculate standardized effect size using SPSS v.20, as appropriate. Power analysis was performed using G*Power.

## Supporting information

not applicable

## List of Supplementary Materials

Materials and Methods

Figure S1. *Ngf* expression in bone marrow of fractured tibiae.

Figure S2. Representative images from calcium imaging using Fura-2 dye.

Figure S3. Adenosine decreases functional activity of mouse DRG neurons.

Figure S4. Adenosine attenuates NGF-induced increase in membrane potential in DRG neurons.

Figure S5. Adenosine mitigates short-term NGF-induced DRG functional activity.

Figure S6. Characterization of PEGDA.

Figure S7. Characterization of A6ACA.

Figure S8. FT-IR spectra of PEGDA-6ACA-PBA macroporous hydrogel and 3-APBA.

Figure S9. ^1^HNMR spectrum of PEGDA-6ACA-PBA macroporous hydrogel.

Figure S10. *In vitro* release of adenosine.

Figure S11. Adenosine levels in circulation of treated mice.

Figure S12. Local delivery of adenosine improves open field activity of fractured animals.

Figure S13. Adenosine receptor gene expression of the whole bone marrow.

## Acknowledgments

We thank Yu-Chih D. Shih for help with statistical analyses and Courtney Karner for providing the *Lepr-cre; tdTomato* mice.

## Author contributions

Conceptualization: YVS, SV

Methodology: YVS, DK, HN, JH, AG

Investigation: YVS, DK

Visualization: YVS, DK, JH

Funding acquisition: SV

Project administration: YVS, SV

Supervision: SV

Writing – original draft: YVS, DK

Writing – review & editing: YVS, DK, SV, DL

## Data and materials availability

All data are available in the main text or the supplementary materials.

