## Supplementary material for "A multi-functional small molecule alleviates fracture pain and promotes bone healing": not applicable

### Supplementary Materials

This PDF file includes:

Materials and Methods

Fig. S1 to S13

References *(82-90)*

**Synthesis and characterization of macroporous hydrogels**

*Synthesis of polyethylene glycol diacrylate (PEGDA):* Poly(ethylene glycol) was acrylated as described elsewhere (*82*). In brief, polyethylene glycol (PEG, MilliporeSigma, Burlington, MA, Cat# P4338) (10 g, ~3 mmol) was dissolved in dry dichloromethane (DCM, 100 mL) at room temperature under argon gas. Triethylamine (MilliporeSigma, Cat# 471283) (627.3 µL, 4.5 mmol) was added to the solution. The reaction mixture was then placed on an ice bath. Acryloyl chloride (MilliporeSigma, Cat# A24109) (364 µL, 4.5 mmol), dissolved in dry DCM (15 mL), was added dropwise to the mixture and the reaction was continued for about 12 h at room temperature. The reaction mixture was then passed through Celite 545 (MilliporeSigma, Cat# 1026931000) and concentrated using a rotary evaporator. The product was precipitated in excess chilled diethyl ether and filtered using Whatman filter paper. The resultant PEGDA was dried overnight under vacuum and purified by using Sephadex G-25 column (GE Healthcare, Chicago, IL) followed by lyophilization. The product was characterized by a combination of FTIR and $^1$HNMR spectroscopy. The FTIR spectra showed peaks at 1725 cm$^{-1}$ corresponding to the ester C=O stretching frequency thus confirming the introduction of acrylate groups in PEGDA *via* ester bond formation (fig. S6A). The diacrylation in PEGDA was further evident from $^1$HNMR as the presence of peaks at 5.81, 6.31 and 6.42 ppm corresponding to vinyl protons of acrylate groups (fig. S6B).

*Synthesis of N-acryloyl-6-aminocaproic acid (A6ACA):* A6ACA was synthesized as described earlier (*83*). 6-Aminocaproic acid (6ACA, MilliporeSigma, Cat# A7824) (13.1 g, 0.1 M) and sodium hydroxide (4.4 g, 0.11 M) was dissolved in 80 mL water. The solution was placed over an ice bath and acryloyl chloride (~10 g, 0.11 M) dissolved in 15 mL dry tetrahydrofuran (THF) was added to the 6ACA solution dropwise. The pH of the reaction mixture was maintained at ~7.8 during the addition of acryloyl chloride by using 2.5 M NaOH solution. After the reaction, the pH of the mixture was gradually decreased to ~3.0 by adding 5 M hydrochloric acid. The product was then extracted using ethyl acetate, concentrated over anhydrous sodium sulfate, and precipitated in chilled n-hexane. The product, A6ACA, was filtered through Whatman filter paper and dried overnight under vacuum at 45 °C. The product was characterized by FTIR and $^1$HNMR spectroscopy. The FTIR spectrum showed peaks at 1655 cm$^{-1}$ and 1543 cm$^{-1}$ corresponding to the amide C=O and N–H stretching frequencies, respectively (fig. S7A). Formation of A6ACA was further evident from $^1$HNMR as the spectrum showed peaks at 5.74, 6.18 and 6.26 ppm corresponding to vinyl protons of acrylamide group (fig. S7B).

*Macroporous hydrogel fabrication:* The macroporous hydrogels containing 3-acrylamido phenylboronic acid (3-APBA, MilliporeSigma, Cat# 771465) (PEGDA-6ACA-PBA) were fabricated using a poly(methyl methacrylate) (PMMA) bead leaching method (*30*). PEGDA (10% w/v), 3-APBA (1 M), A6ACA (0.5 M) and Irgacure 2959 (MilliporeSigma, Cat# 410896, 0.5% w/v) were dissolved in 20:80 water:ethanol mixture. 20 μL of the mixture was added into a cylindrical polypropylene mold (~5 mm in diameter) packed with ~20 mg of PMMA microbeads (150-180 μm, Cospheric, Santa Barbera, CA). The mixture was photopolymerized by UV light irradiation for about 10 min. The resulting hydrogel network embedded with the PMMA beads was incubated in acetone for 3 days to remove the beads with frequent changes of solvent. The macroporous hydrogel was washed and rehydrated with deionized water. FTIR spectrum of the freeze-dried hydrogel showed a peak at 1725 cm$^{-1}$ corresponding to the ester C=O stretching frequency indicating the presence of PEGDA (fig. S8). Presence of phenylboronic acid (PBA) moieties was confirmed *via* $^1$HNMR spectroscopy. For NMR measurements, freshly prepared porous hydrogels were thoroughly washed with DI water to remove unreacted precursors and then freeze-dried. The freeze-dried samples were minced and fully suspended in $D_2O$ by adding 5 M NaOH solution in $D_2O$. The $^1$HNMR spectrum, recorded using a 500 MHz Varian spectrometer, showed peaks at 7.07-7.27 ppm corresponding to aromatic protons of PBA (fig. S9). The PBA content in the hydrogel was quantified by using UV/Vis spectrophotometer. Towards this, the macroporous hydrogels were completely dissociated in 5 M NaOH solution in water. The PBA content in the solution was determined by using a UV/Vis spectrophotometer at an absorption wavelength of 255 nm. A standard calibration curve of absorbance *vs* concentration was generated using free 3-APBA solutions of known concentrations (21.25 to 170 μg/mL) and used to calculate PBA content. The estimated value suggest that the PBA constituted over 45% of the dry weight of hydrogels which further indicated that almost 93±3% of 3-APBA was incorporated into the hydrogel network. Macroporous hydrogels without 3-APBA (PEGDA-6ACA) was prepared similarly and used as controls for animal studies. For sterilization, the hydrogels were soaked in 70% ethanol for 6 h and washed extensively in phosphate buffered saline (PBS) for 3-4 days.

*Adenosine loading:* To load adenosine (MilliporeSigma, Cat# A4036) into the macroporous hydrogel (PEGDA-6ACA-PBA), hydrogel discs were incubated in adenosine solution in PBS at 6 mg/mL for about 6 h at 37 °C and washed thoroughly to remove any unbound adenosine. To measure the amount of adenosine loaded, the discs were soaked in acetate buffer (0.1 M, pH 3.5) for about 2 h to release the adenosine into the buffer. The adenosine content in the buffer was determined by using a UV/Vis spectrophotometer at wavelength of 260 nm. A standard calibration curve of absorbance *vs* concentration was generated using adenosine solutions of known concentrations (3.9-125 µg/mL) and used to calculate the adenosine concentration.

*Adenosine release:* To determine the release profile of adenosine, adenosine loaded macroporous hydrogels were incubated in α-MEM (ThermoFisher Scientific, Waltham, MA, Cat# 12561056) containing 10% (v/v) fetal bovine serum (FBS) at 37 °C. At predetermined time intervals, 20 vol% of the medium was removed and supplemented with fresh medium. The concentration of adenosine in the released medium was determined *via* UV/Vis absorption spectroscopy using a standard calibration curve as described above.

**Behavioral tests**

*Weight bearing*

Static weight bearing was measured using an incapacitance meter (IITC Life Science, Woodland Hills, CA). Prior to measurements, mice were trained on the incapacitance meter for 5 days (10 min/day). Data were collected if the animal stood in an upright position and facing front, without noticeable weight shift, lifting or offloading a limb, or turning the head. Each animal was tested for 5 to 6 trials and the weight of both hind limbs was recorded. Weight bearing of the fractured limb (i.e., ipsilateral, right) was expressed as a percentage of the total weight borne by the hindlimbs calculated by: (weight of ipsilateral hindlimb/total weight of both hindlimbs)*100.

*Open Field Activity*

Open field activity was performed at Duke Mouse Behavioral and Neuroendocrine Core similar to ref. (*84*). Mice were acclimated to the room for a day prior to testing. Mice were placed individually into the VersaMax open field activity monitoring system with clear acrylic test chambers (40 X 40 X 30 cm) and a grid of infrared photobeams containing photocells and sensors to collect and analyze vertical, horizontal, and stereotypic activity (AccuScan

Instruments, Columbus, OH). Locomotion was monitored over 60 min at 5 min intervals. The vertical activity (units), vertical movement time (s), ambulatory activity count (units), ambulatory time (s), total distance traveled (cm), and rest time (s) were determined by the software. Results were calculated as a ratio of post-fracture divided by pre-fracture (baseline) values.

**Histology**

For safranin O staining, tibiae were fixed with 4% PFA at 4°C for 1 day and decalcified using 14% ethylenediaminetetracetic acid (EDTA, pH 7.3) for 2 weeks at 4°C with constant shaking. The samples were gradually dehydrated using increasing concentrations of ethanol and incubated in Citrisolv (Decon Laboratories, King of Prussia, PA) until equilibrium was reached. Following dehydration, samples were immersed in a mixture of 50% (v/v) Citrisolv and 50% (w/w) paraffin (General Data Healthcare, Cinicinnati, OH) for 30 min at 70 °C. The samples were embedded in paraffin and 7-µm thick sections were generated using a rotary microtome (Leica Microsystems, Buffalo Grove, IL, RM2255). For immunofluorescence staining of DRG frozen sections, L3-L4 DRG were dissected, fixed with 4% PFA at 4°C for 2 h, and incubated in 30% sucrose overnight, embedded in OCT, and 5-µm thick sections were generated by using a cryostat (Leica, CM1850).

**MicroCT**

Tibiae were collected, fixed in 4% PFA at 4 °C for 1 day, and rinsed with PBS. The fixed samples were placed in 50-mL centrifuge tubes with styrofoam spacers and loaded into a microcomputed tomography (microCT) scanner (vivaCT 80, Scanco Medical, Wayne, PA). The samples were scanned at 55 keV at a pixel resolution of 10.4 µm. The reconstruction of the images was performed using microCT Evaluation Program V6.6 (Scanco Medical), followed by generation of radiographs and 3D models using microCT Ray V4.0 (Scanco Medical). Total volume (TV), bone volume (BV) per total volume (TV) (%BV/TV), and bone mineral density (BMD) was quantified using the phantom as a reference based on 100 contiguous slices.

**Cell isolation and *in vitro* culture**

*DRG neurons:* DRG were isolated from mice using a modified version of the previously described protocol (*85*). Briefly, mice were euthanized and dosed in 70% ethanol to prevent fur contamination. Mice were positioned prone, and an incision was made along the spine from the neck to the base of the tail. Under a dissecting microscope, the surrounding soft tissue of the spinal column was removed from the dorsal side by Friedman-Pearson Rongeurs (Fine Science Tools, Foster City, CA, Cat# 16021-14) from the mid thoracic region to the lumbosacral joint. The vertebral bone surrounding the DRG was gently crushed and removed with rongeurs exposing the spinal cord. The DRG were carefully extracted near the foramen between the vertebral levels. L3 and L4 DRG in fractured animals were collected for analyses. For other experiments DRG (L1-L5) were collected. Dissected DRG were placed directly into ice-cold DMEM-F12 medium (ThermoFisher, Cat# 11320033) for culture, Trizol (ThermoFisher, Cat# 15596018) for PCR, or 4% PFA for fixation. For cell culture experiments, collected DRG were digested in digestion solution comprised of collagenase type II (Worthington, Lakewood, NJ, Cat# LS004176) and dispase (1.5 mg/mL each) dissolved in DMEM/F12. DRG were agitated in a digestion solution on an orbital shaker at 60 rpm and 37°C for 20 minutes and repeated thrice with fresh digestion solution. Next, the digestion solution was replaced with trypsin-EDTA (0.025%) in DMEM/F12 and incubated for an additional 15 min to disrupt the remaining cell-to-cell adhesions. The solution was then replaced with DMEM/F12 and FBS (1:3) to neutralize trypsin. DRG were triturated to create a cell suspension and gently layered onto 15% bovine serum albumin (BSA) solution without mixing and centrifuged for 6 min at 280 g with minimal acceleration and no deceleration to separate nonneuronal cells and debris (*86*). The supernatant was carefully aspirated, and the sensory neurons were resuspended in DRG culture medium. To culture DRG neurons, culture media composed of Neurobasal-A medium (ThermoFisher, Cat# 10888022), 2% B27 supplement (ThermoFisher, Cat# 17504044), 1% glutamax (ThermoFisher, Cat# 35050061), and penicillin/streptomycin (10000 U/mL, 1% v/v, ThermoFisher, Cat# 15140122) was used. Cells were treated with or without NGF (200 ng/mL) for 24 h prior to immunofluorescence imaging. Cells were exposed to NGF (200 ng/mL) for 24 h (long-term) for calcium and FluoVolt imaging and 1 h (short-term) for calcium imaging. Cells were cultured on custom-made glass surface with silicone wells. Briefly, cover slides (#1 thickness) were cleaned by agitating in 0.5 M NaOH for 30 minutes followed by subsequent rinsing in $dH_2O$ and 100% ethanol and air-dried. Wells were produced in cured polydimethylsiloxane (PDMS, Sylgard 184,

Ellsworth Adhesives, Germantown, WI) with a 8-mm biopsy punch and bonding to the cover slides. The wells were coated with poly-lysine by treating overnight with 0.1 mg/mL poly-d-lysine solution, rinsed with sterile water and air-dried. These custom-fabricated wells were stored for up to 7 days prior to use. The wells were treated with laminin (20 µg/mL) for at least 4 hours prior to cell seeding. Laminin solution was aspirated out, incubated with culture medium for 1 hour prior to culturing DRG neurons.

*MSC:* MSCs were isolated as previously described with some modifications (*87*). Briefly, the femurs, tibiae, vertebrae of mice were harvested, crushed with pestle and mortar in harvest buffer 1% v/v FBS in PBS to release bone marrow (BM) tissue, filtered through a 40-µm cell strainer, and centrifuged at 200 rcf. Cells were seeded in a 24-well plate at a cell density of 1 million cells/cm$^2$ in growth media (GM) containing α-MEM, FBS (10% v/v, ThermoFisher, Cat# 16000044), penicillin/streptomycin (10000 U/mL; 1% v/v) in cultured in humidified incubator (37°C, 5% $CO_2$). The medium was replaced after 3 days and further cultured for 6 days before passage. For passaging, cells were incubated in 0.25% trypsin-EDTA for 2 min at 37°C, detached with a cell scraper, neutralized by using GM, centrifuged, and sub-cultured at a density of 8000 cells/cm$^2$. All experiments were performed between 1-2 passage. Osteogenic medium (OM) was prepared by supplementing GM with 10 mM β-glycerophosphate (MilliporeSigma, Cat# G9422), 50 µM ascorbic acid-2-phosphate (MilliporeSigma, Cat# A8960), and 100 nM dexamethasone (MilliporeSigma, Cat# D4902).

**Calcium imaging**

Intracellular cytosolic $Ca^{2+}$ was evaluated by Fura-2 loaded DRG neurons (*88*). Fura-2 (ThermoFisher, Cat# F1221) stock aliquots (2 mM in 100% DMSO) were mixed 1:1 in Pluronic F-127 and then 1:500 in Tyrode's solution (140 mM NaCl, 5 mM KCl, 2 mM $CaCl_2$, 2 mM $MgCl_2$, 10 mM HEPES, 10 mM glucose, pH 7.4) (*88*). DRG samples were washed twice in Tyrode's solution before replacement with Fura-2 loading solution and incubated at room temperature for 40 min. Samples were then washed thrice with Tyrode's solution and incubated for an additional 20 min prior to loading for imaging. Cells were mounted on the translation stage of an Olympus IX81 inverted microscope (Olympus America, Center Valley, PA) and depending on the experimental group, were treated with either adenosine (5 µM), or adenosine (5 μM) along with DPCPX (100 nM) for 5 min before stimulation with capsaicin (100 nM). For the experiments involving DPCPX, since DPCPX was dissolved in DMSO all corresponding control cultures were exposed to the vehicle DMSO.

Fura-2 dual excitation and emission was accomplished using 340- and 380-nm excitation filters and a 510 nm emission filter, and cells were visualized with an Olympus UPlan FLN 20X 1.3 NA water immersion objective. Light was supplied by a Lambda XL (Sutter Instrument Company, Novato, CA) using variable aperture. Digital images (150-ms exposure) were recorded with a Hamamatsu EM CCD camera (Hamamatsu Photonics, Hamamatsu City, Japan) at 1 s intervals. Imaging was performed by first establishing a baseline intensity ratio (340/380 nm) for the region of interest (ROI) prior to capsaicin (TRPV1 agonist) stimulation. The normalized Fura-2 intensity profiles were plotted as the ratiometric intensity divided by the baseline intensity. Peak intensity measurements are reported as the maximum normalized intensity during stimulation. The total number of DRG neurons in a given ROI was determined by adding KCl at the end of the experiment and counting the number of activated DRG neurons, i.e., cells with a ratiometric change.

**Membrane potential imaging**

DRG neuron potential changes were evaluated with FluoVolt, a voltage sensitive indicator dye (ThermoFisher, Cat# F10488). Fluovolt was loaded according to manufacturer instructions; briefly, loading solution was prepared by adding 10 μL of 10X component B and 1 μL of component A in a 1.5-mL tube, followed by 1 mL of Tyrode's solution. DRG culture medium was removed and washed prior to adding the loading solution, followed by incubating at room temperature for 30 min. Loading solution was then removed and washed thrice with Tyrode's solution. Imaging was performed on translation stage of an Olympus IX81 inverted microscope (Olympus America, Center Valley, PA, USA, with FITC excitation filter. Digital images (250-ms exposure) were recorded with a Hamamatsu EM CCD camera (Hamamatsu Photonics) at 1 s intervals. Imaging was performed prior to capsaicin stimulation to determine the baseline intensity for the DRG neurons. Fluorescence intensity profiles were reported as the intensity at a given time point divided by the baseline intensity. Peak intensity measurements are reported as the maximum normalized intensity during stimulation.

**RT-qPCR**

Cells or tissues were analyzed for gene expression by quantitative real-time polymerase chain reaction (RT-qPCR). Nucleic acids were extracted with TRIzol, phase-separated with chloroform, and precipitated in isopropanol. One microgram of RNA was reverse transcribed using iScript cDNA Synthesis Kit (Bio-Rad, Hercules, CA, Cat# 1708891), according to the manufacturer's instructions (*89*). Quantitative PCR was performed with iTaq Universal SYBR green reagent (Bio-Rad, Cat# 1725124) with denaturation at 95 °C for 30 s for one cycle, and amplification (denaturation + annealing/extension) at 95 °C for 5 s and 60 °C for 30 s for 40 cycles on a polymerase chain reaction (PCR) cycler (Bio-Rad, CFX96 Touch). The primer sequences used are: *Ngf* (forward, GGGAG CGCAT CGAGT TTTG; reverse, TACGC TATGC ACCTC ACTGC), *Trpv1* (forward, CAGCG AGTTC AAAGA CCCAG A; reverse, GCAGA GCAAT GGTGT CGTTC), *Adora1* (forward, CCCCA TCGTCTA TGCCTTCC; reverse, CATCG GAAGT GGTCG TTCCA) , *Adora2a* (forward, GCCAG AGCAA GAGGC AGGTA T; reverse, TCCCA AAGGC TTTCT CACGG), *Adora2b* (forward, ATCTT TAGCC TCTTG GCGGT G; reverse, GACCC AGAGG ACAGC AATGA T), *Adora3* (forward, GCTGTA GACCGA TACCTG CG; reverse, GGAAAC TAGCCA GCAAAG GC), *Runx2* (forward, TGGCC GGGAA TGATG AGAAC; reverse, TGAAA CTCTT GCCTC GTCCG), *Sp7* (forward, TGCCT GACTC CTTGG GACC; reverse, TAGTG AGCTT CTTCC TCAAG CA), *Ibsp* (forward, TCCAC ACTTT CCACA CTCTC G; reverse, CTTTC TGCAT CTCCA GCCTT C), *Bglap* (forward, GCTAC CTTGG AGCCT CAGTC; reverse, AGGGT TAAGC TCACA CTGCT C), *18S ribosomal RNA* (forward, ACCAG AGCGA AAGCA TTTGC CA; reverse, ATCGC CAGTC GGCAT CGTTT AT), *Gapdh* (forward, GCACA GTCAA GGCCG AGAAT; reverse, GCCTT CTCCA TGGTG GTGAA) [14]. The expression level of each target gene was normalized to the housekeeping gene and to their respective controls and presented as fold change expressed as $2^{-\Delta\Delta Ct}$ values. *NGF* expression was normalized to *Gapdh*, and other gene expressions were normalized to *18S rRNA*.

**Immunofluorescence staining**

*In vitro* cultured DRG neurons, frozen DRG sections, and paraffin-embedded tibiae sections were stained and imaged. *In vitro cultured DRG neurons:* cells were fixed in 4% PFA for 15 min, permeabilized in 0.1% Triton X-100 in PBS for 10 min, followed by blocking for 1 h in 3% BSA in PBS at room temperature. Cells were co-stained with TUBB3 (Novus Biologicals, Littleton, CO, Cat# NB100-1612, 1:500 dilution in 3% BSA) and TRPV1 (Novus Biologicals, Cat# NBP1-71774, 1:300 dilution in 3% BSA) antibodies at 4°C overnight. Cells were washed with PBS at room temperature for 10 min thrice, and incubated with secondary antibodies donkey anti-chicken AlexaFluor 488 (Jackson ImmunoResearch, West Grove, PA, Cat# 703-545-155, 1:300 dilution) and donkey anti-rabbit AlexaFluor 647 (Jackson ImmunoResearch, Cat# 711-605-152, 1:300) for 1 h at room temperature. *Frozen DRG sections:* sections were permeabilized in 0.1% Triton X-100 in PBS for 10 min, followed by blocking for 1 hour in blocking solution comprised of 3% BSA, 0.26 M glycine, 5% normal donkey serum in Tris buffered saline (TBS) at room temperature. Sections were co-stained with TUBB3 (Novus Biologicals, Cat# NB100-1612, 1:500 dilution in blocking solution) and TRPV1 (Novus Biologicals, Cat# NBP1-71774, 1:300 dilution in blocking solution) antibodies, or co-stained with TUBB3 (Novus Biologicals, Cat# NB100-1612, 1:500 dilution in blocking solution) and ADORA1 (Proteintech, Rosemont, IL, Cat# 55026-1-AP, 1:100 dilution in blocking solution) antibodies at 4°C overnight. Sections were washed with TBS with 0.1% Tween-20 (TBS-T) at room temperature for 10 min thrice, and then incubated with secondary antibodies donkey anti-chicken AlexaFluor 488 (Jackson ImmunoResearch, Cat# 703-545-155, 1:300 dilution in blocking solution) and donkey anti-rabbit AlexaFluor 647 (Jackson ImmunoResearch, Cat# 711-605-152, 1:300 dilution in blocking solution) for 1 h at room temperature. *Paraffin-embedded tibiae sections:* sections were heated for antigen retrieval in citrate buffer (MilliporeSigma, Cat# C9999) for 20 min, permeabilized in 0.1% Triton X-100 in PBS for 10 min, followed by blocking for 1 h in blocking solution comprised of 3% BSA, 0.26 M glycine, 5% normal donkey serum in Tris buffered saline (TBS) at room temperature. Sections were co-stained for Td-tomato (MyBiosource, San Diego, CA, Cat# MBS448092 1:100 dilution in blocking solution) and ADORA2B (MyBiosource, Cat# MBS8207549, 1:100 dilution in blocking solution) antibodies at 4°C overnight. Sections were washed with TBS-T at room temperature for 10 min thrice, and then incubated with secondary antibodies donkey anti-goat FITC (Jackson ImmunoResearch, Cat# 705-096-147) and donkey anti-rabbit AlexaFluor 647 (Jackson ImmunoResearch, Cat# 711-605-152, 1:300) for 1 h at room temperature. Finally, all samples were washed with PBS or TBST, covered with mounting solution (ThermoFisher, Cat# P36971), sealed and imaged using

Cy5 and GFP filters on a Keyence BZ-X700 microscope. Fluorescence intensity was quantified by ImageJ software and presented in arbitrary units.

**Safranin O staining**

To visualize the callus remodeling, tissue sections were stained with 1% safranin-O (Millipore Sigma, Cat# S8884) for 1 h and counter-stained with 0.02% Fast Green (Millipore Sigma, Cat# F7258) for 1 min. The sections were then dehydrated in an ethanol gradient, mounted with Cytoseal (ThermoScientific, Cat# 23-244256), and imaged by using a Keyence BZ-X710 microscope.

**Measurement of adenosine in plasma**

Peripheral blood was collected and immediately incubated in ice cold stop solution at a 1:2 ratio (blood to stop solution) to inhibit degradation of adenosine (*90*). Stop solution comprised of 0.2 mM dipyridamole (Tocris, Minneapolis, MN), 5 µM erythro-9(2-hydroxy-3-nonyl)-adenine (EHNA; Tocris), 62 µM Adenosine 5'-(α,β-methylene)diphosphate sodium salt (APCP), 5 mM EDTA, and 25 IU/mL heparin (Tocris) in PBS. The blood was centrifuged at 2000 rcf for 10 min at 4°C to separate cells from plasma. Adenosine assay kit (Cell Biolabs, San Diego, CA, Cat# MET-5090) was used to measure adenosine levels from plasma according to manufacturer's protocol. Briefly, a reaction mixture comprised of fluorometric Probe, HRP, adenosine deaminase, purine nucleoside phosphorylase, xanthine oxidase, assay buffer and a control mix comprised of fluorometric Probe, HRP, purine nucleoside phosphorylase, xanthine oxidase, assay buffer were made and mixed with 50 µL of plasma for 15 min. at room temp. The relative fluorescence unit (RFU) was measured using a microplate reader with excitation at 570 nm and emission at 590 nm. Adenosine concentration was determined using a standard curve of known concentrations, and subtracted for the background by using the values in control mix.

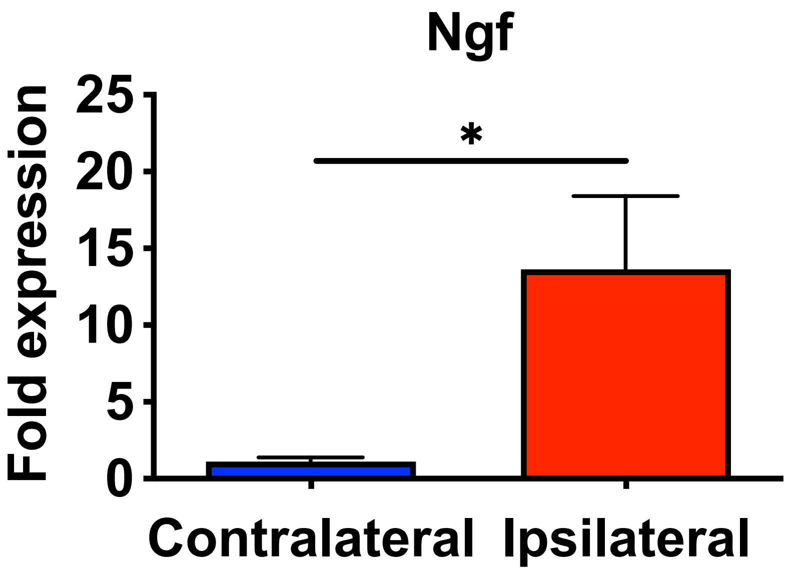


**Figure S1.** ***Ngf*** **expression in bone marrow of fractured tibiae.** Relative nerve growth factor (*Ngf*) gene expression between whole bone marrow of contralateral and ipsilateral tibiae of fractured mice at 5 days post-fracture (mean ± SEM, $n$=3 mice per group. Two-tailed unpaired $t$ test). *$P < 0.05$.

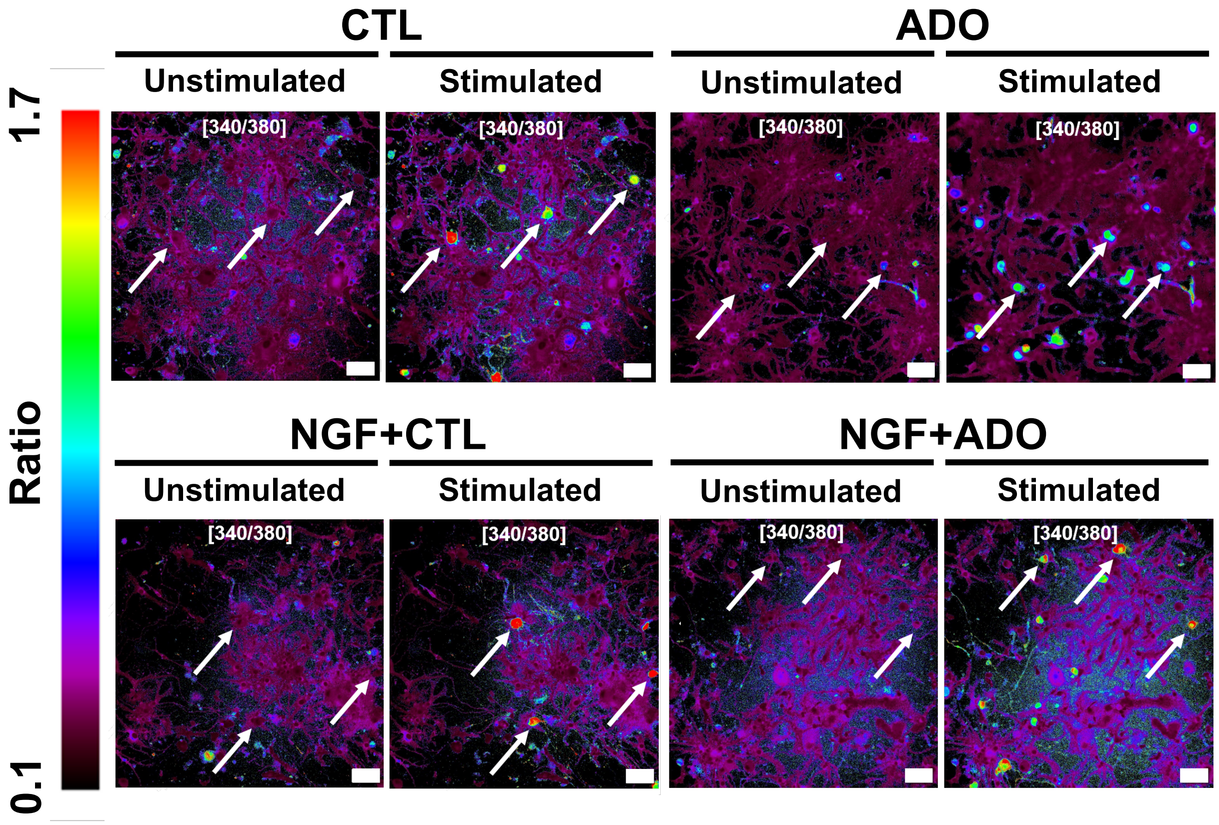


**Figure S2. Representative images from calcium imaging using Fura-2 dye.** TRPV1 agonist capsaicin was used to stimulate the cells and signals were normalized to the baseline (unstimulated, *n*=8-16 cells per group). Arrows indicate cells with changing fluorescence after stimulation with TRPV1 agonist capsaicin.

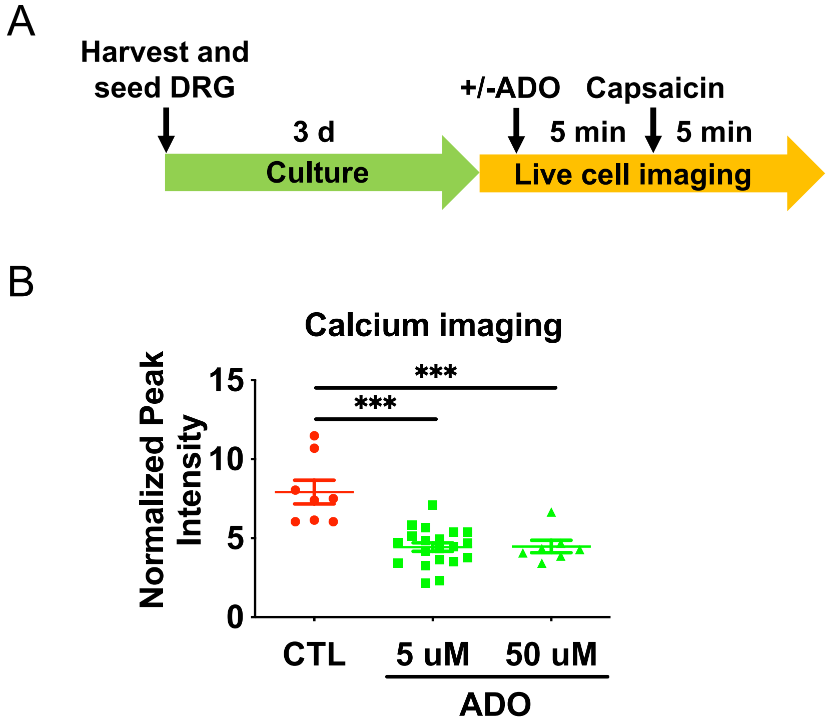


**Figure S3. Adenosine decreases functional activity of mouse DRG neurons.** (**A**) Schematic of calcium imaging experiment. (**B**) Normalized peak intensity of dissociated dorsal root ganglion (DRG) neurons *in vitro* treated with two different concentrations of adenosine (ADO) and stimulated by TRPV1 agonist capsaicin (mean ± SEM, $n$=7-18 cells per group. Two-tailed unpaired $t$ test). ***$P < 0.001$.

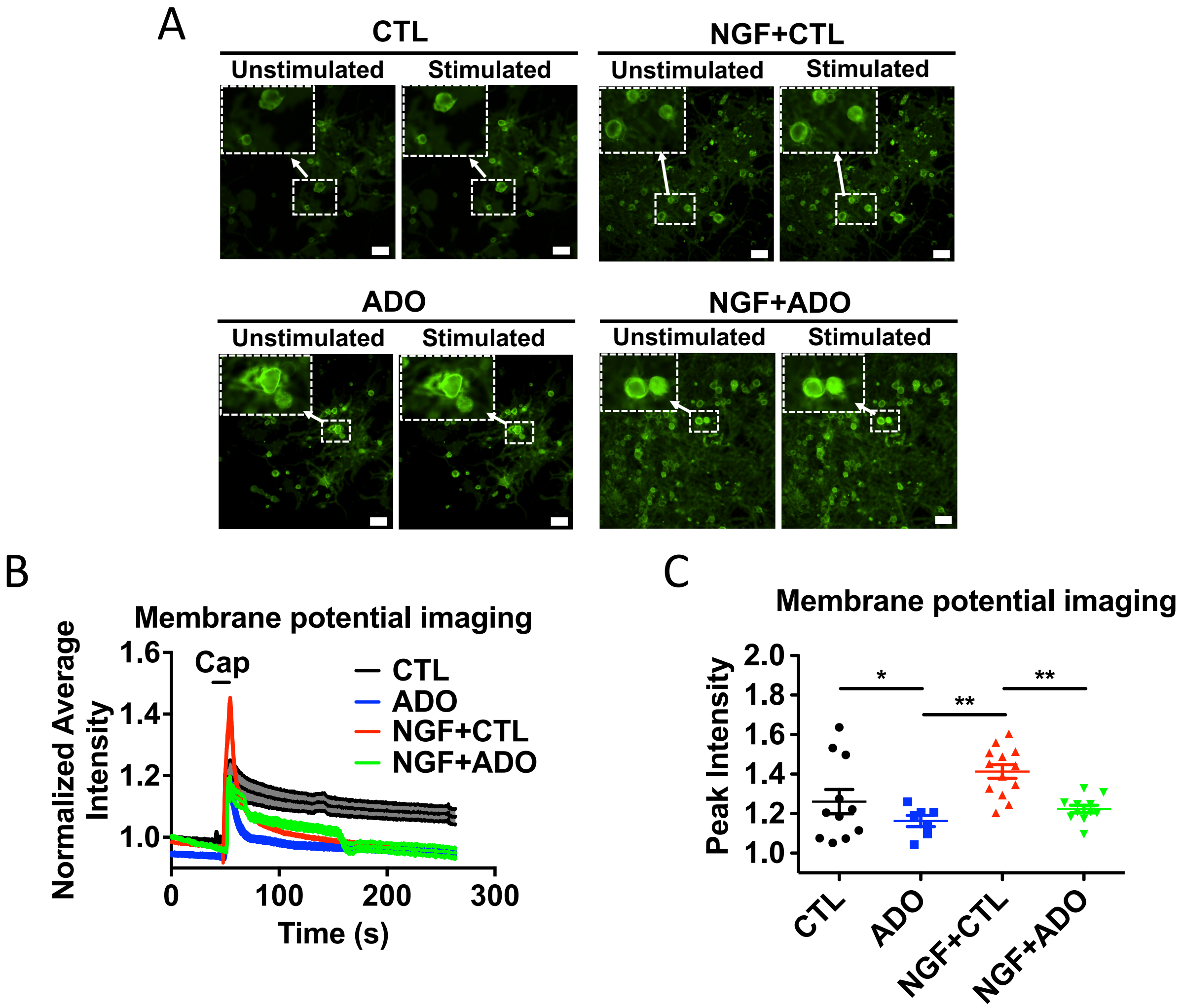


**Figure S4. Adenosine attenuates NGF-induced increase in membrane potential in DRG neurons.** (**A**) Relative fluorescence intensity of membrane potential imaging of dissociated DRG neurons after 1 d of NGF treatment followed by adenosine. Magnified views indicate cells with changing fluorescence intensity after stimulation with TRPV1 agonist capsaicin (Cap; stimulated) was added at the specified time (black line). (**B**) Normalized average signal intensity from membrane potential imaging. (**C**) Normalized peak intensity from membrane potential imaging (mean ± SEM, *n*=7-13 cells per group. Two-way ANOVA with Tukey post hoc test). $*P<0.05$, $**P < 0.01$.

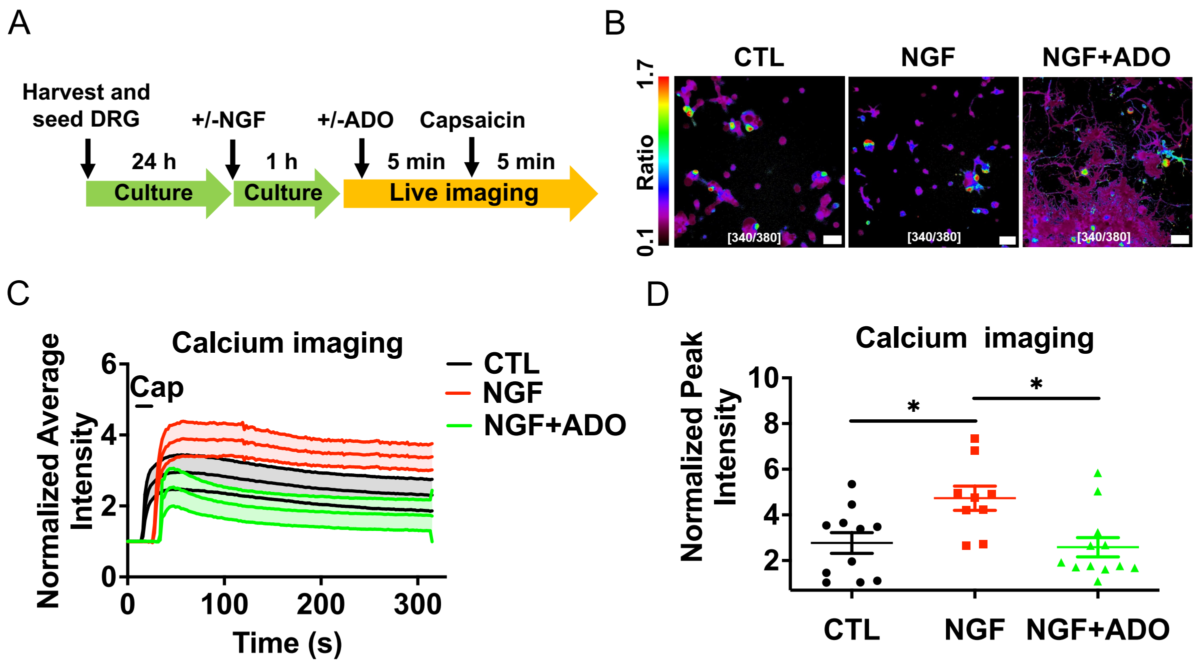


**Figure S5. Adenosine mitigates short-term NGF-induced DRG functional activity. A**) Experimental design to probe the effect of 1 h of nerve growth factor (NGF) treatment followed by adenosine on activity of dissociated dorsal root ganglion (DRG) neurons using calcium imaging. (**B**) Representative ratiometric images from calcium imaging using Fura-2 dye. TRPV1 agonist capsaicin was used to stimulate cells and signals were normalized to baseline. (**C**) Normalized average signal intensity from the calcium imaging. TRPV1 agonist capsaicin (Cap) was added at the specified time (black line) (**D**) Normalized peak intensity from the calcium imaging (mean ± SEM, $n$=9-12 cells per group. Two-way ANOVA with Tukey post hoc test). *$P$<0.05, **$P$ < 0.01.

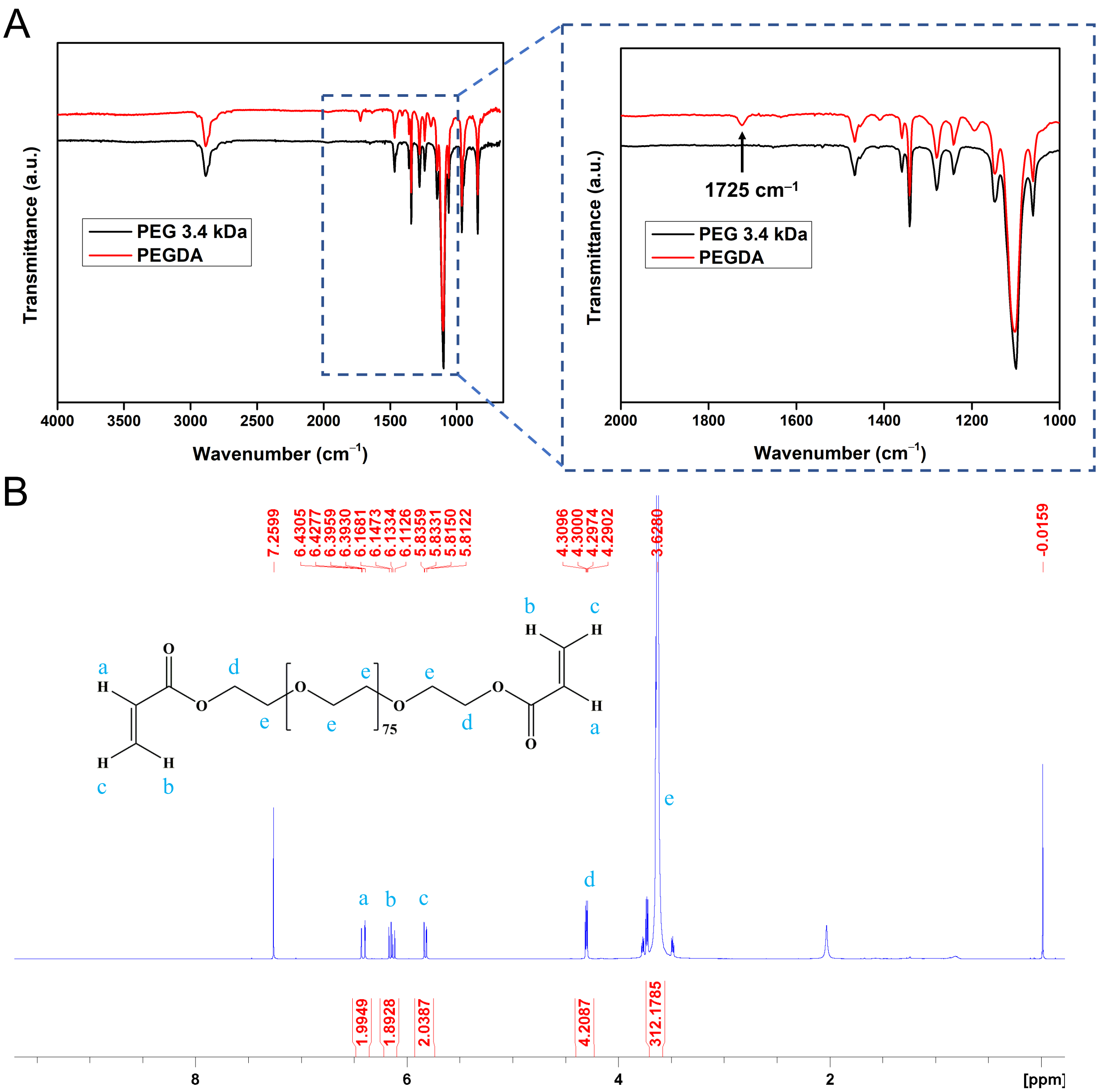


**Figure S6. Characterization of PEGDA.** (A) FTIR and (B) $^1$HNMR spectrum of polyethylene glycol diacrylate (PEGDA). FTIR spectra were recorded using ZnSe crystal in attenuated total reflectance (ATR) mode. Arrow at 1725 cm$^{-1}$ indicates the stretching frequency for the ester C=O bond in PEGDA. $^1$HNMR spectrum of PEGDA recorded in $CDCl_3$ at 25 °C.

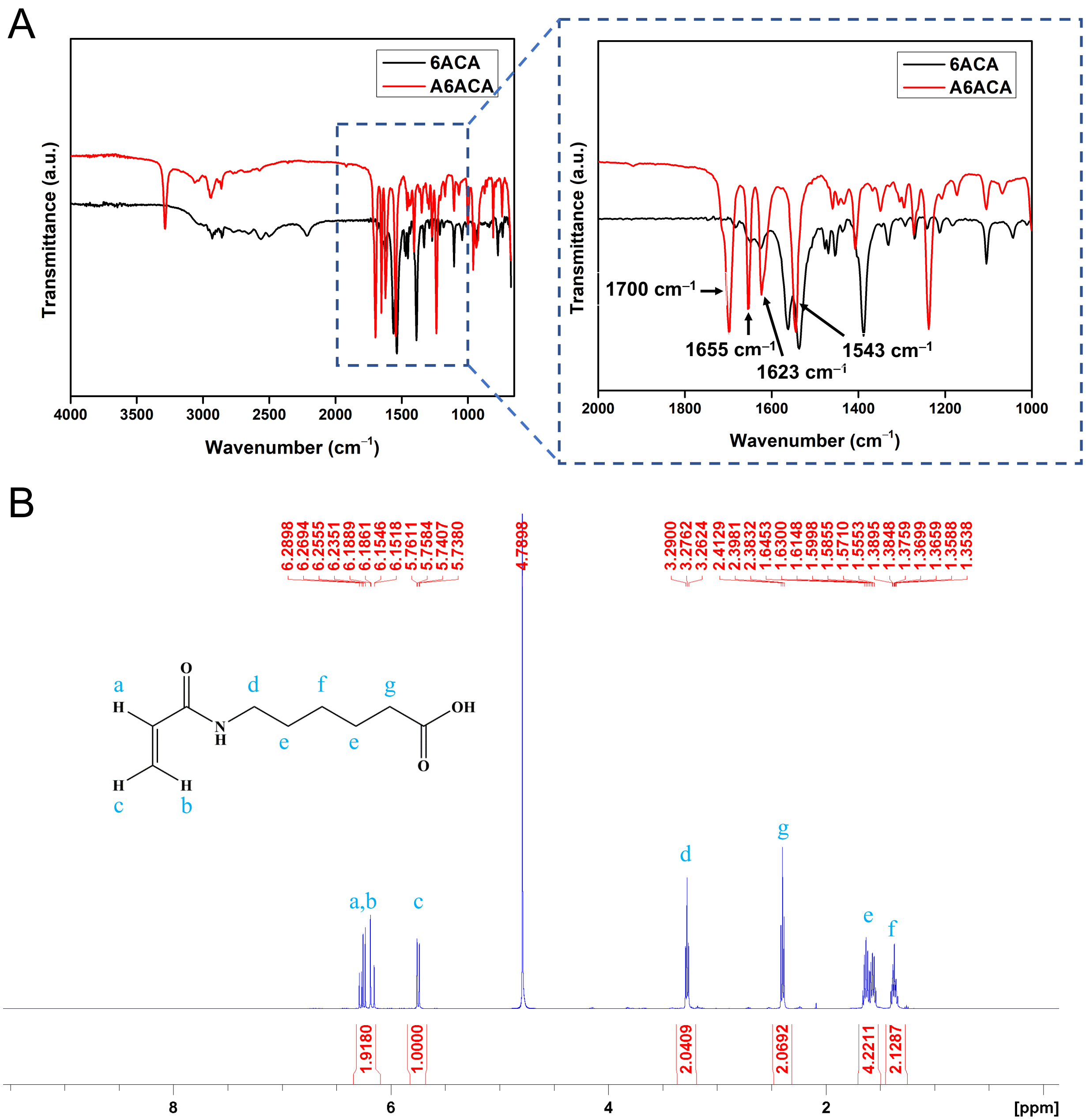


**Figure S7. Characterization of A6ACA.** (A) FTIR and (B) $^{1}$HNMR spectrum of N-acryloyl-6-aminocaproic acid (A6ACA). FTIR spectra were recorded using ZnSe crystal in attenuated total reflectance (ATR) mode. Arrows at 1655 cm$^{-1}$ and 1543 cm$^{-1}$ corresponding to the amide C=O and N–H stretching frequencies, respectively. $^{1}$HNMR spectrum of A6ACA recorded in $D_2O$ at 25 °C.

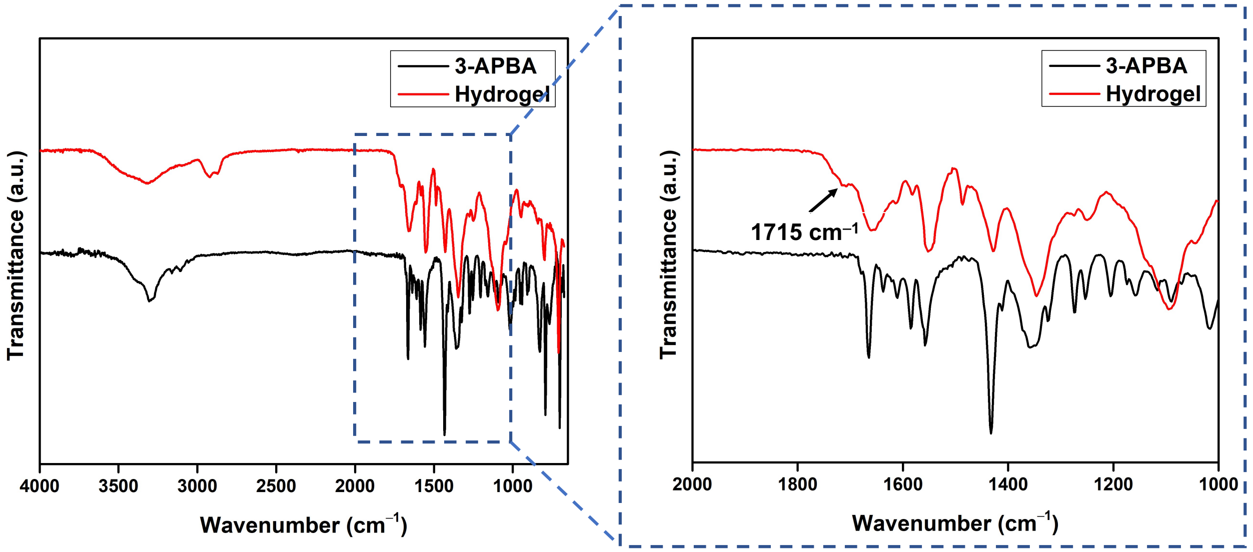


**Figure S8. Characterization of PEG-6ACA-PBA macroporous hydrogel.** FTIR spectra of the lyophilized gel recorded using ZnSe crystal in attenuated total reflectance (ATR) mode. Arrow at 1715 $cm^{-1}$ indicates the stretching frequency for the ester C=O bond of PEGDA in PEG-6ACA-PBA hydrogel.

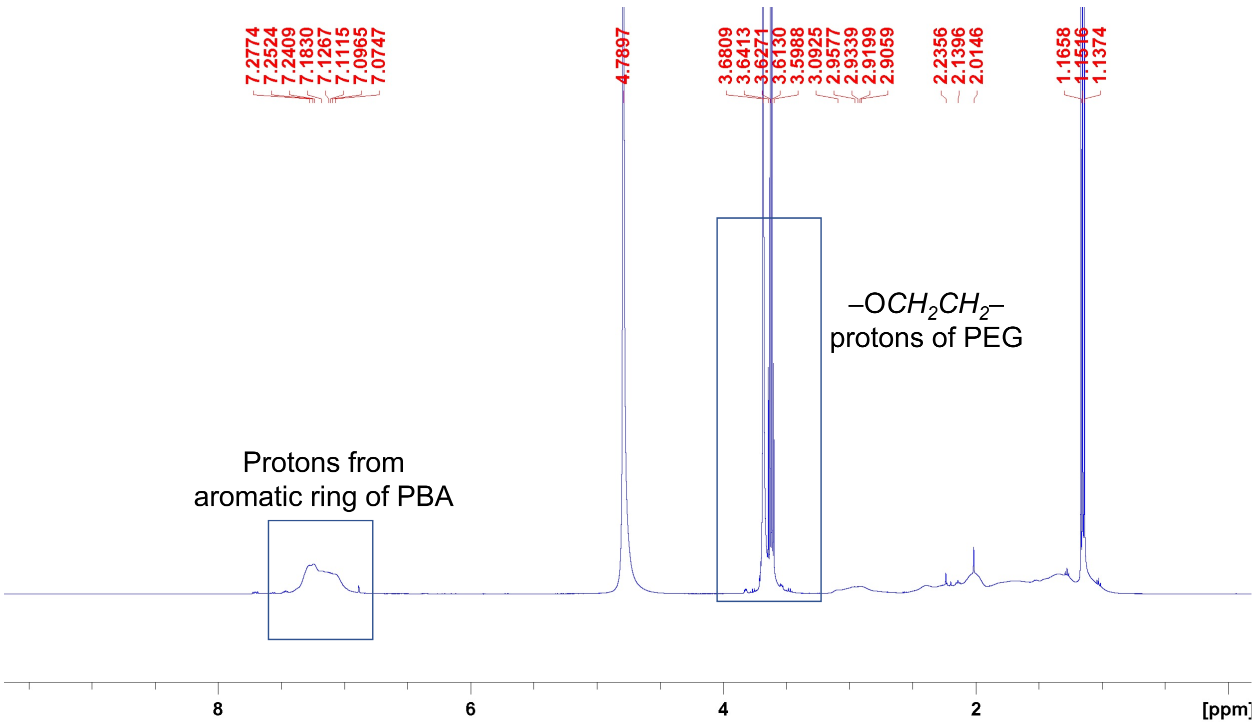


**Figure S9. Characterization of PEG-6ACA-PBA macroporous hydrogel.** $^{1}$HNMR spectrum of the hydrogel recorded in $D_2O$ at 25 °C.

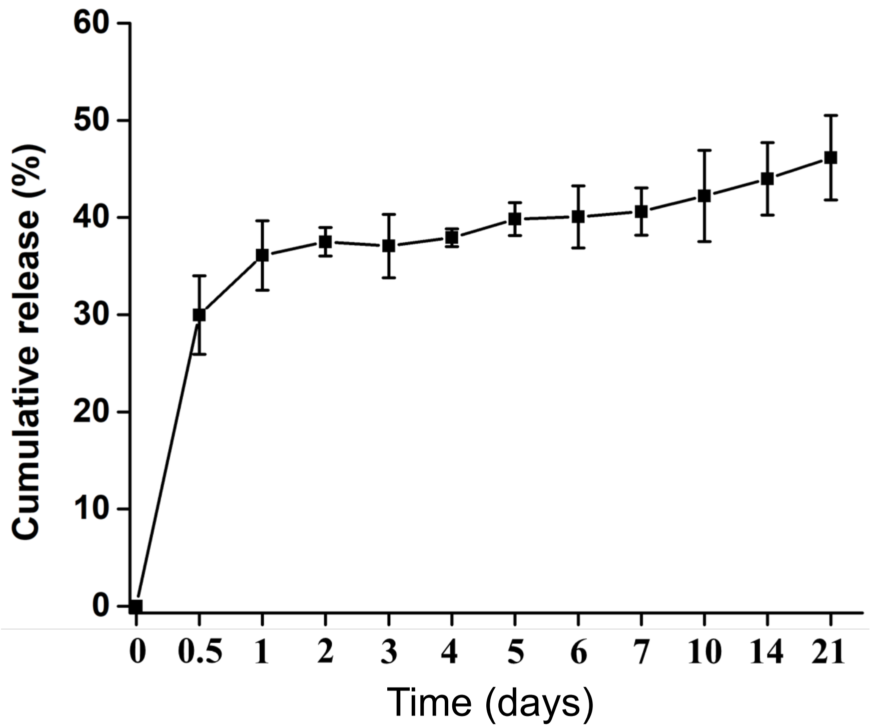


**Figure S10. *In vitro* release of adenosine.** Cumulative percentage of *in vitro* release of adenosine from PEGDA-6ACA-PBA macroporous hydrogel over 21 days (*n*=3).

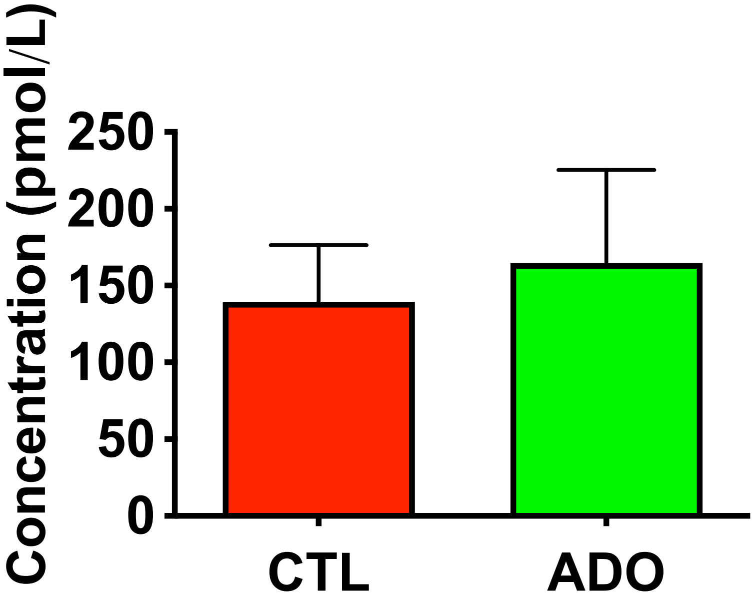


**Figure S11. Adenosine levels in circulation of treated mice.** Concentration of adenosine in peripheral blood of fractured mice treated with control or adenosine-loaded macroporous hydrogel at 3 days post fracture (mean ± SEM, *n*=6 measurements from 3 mice. Mann Whitney U test).

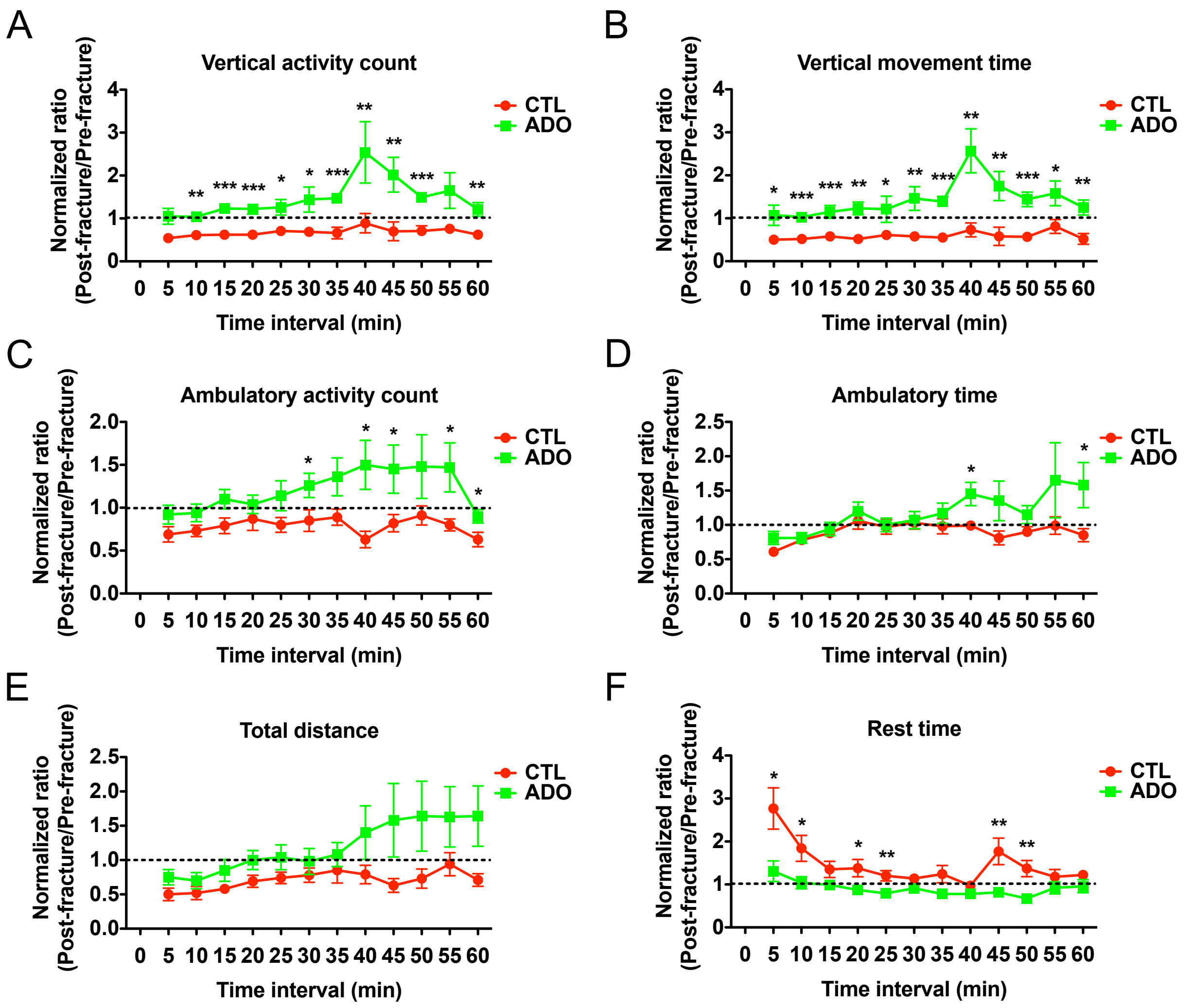


**Figure S12. Local delivery of adenosine improves open field activity of fractured animals.** (**A**) Normalized ratio of vertical activity count, (**B**) vertical movement time, (**C**) ambulatory activity count, (**D**) ambulatory time, (**E**) total distance, and (**F**) rest time of treated mice at 5-min intervals at 7 days post fracture. Results are normalized to the pre-fracture values (mean ± SEM, $n$=9 mice per group. Mann Whitney U test). *$P$<0.05, **$P$ < 0.01, ***$P$ < 0.001.

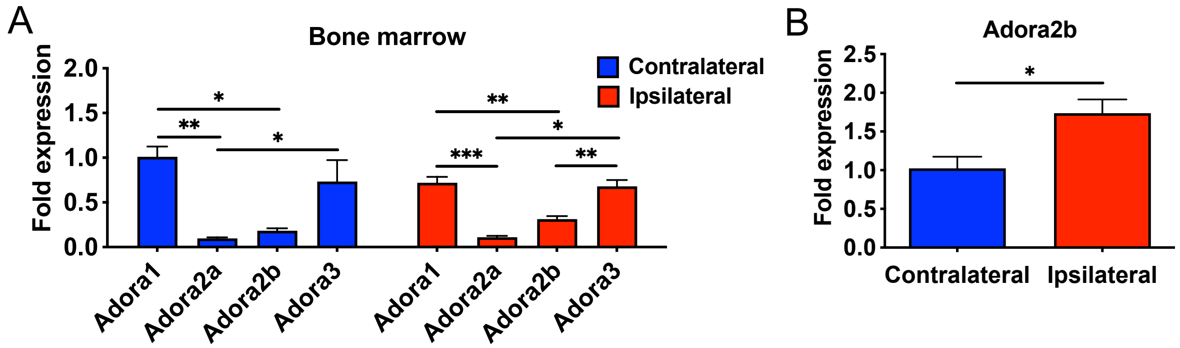


**Figure S13. Adenosine receptor gene expression of the whole bone marrow.** (**A**) Relative gene expression of adenosine receptors in whole bone marrow of fractured mice (mean ± SEM, *n*=3 mice per group. One-way ANOVA with Tukey post hoc test was used for statistical analysis. (**B**) Relative gene expression of *Adora2b* of the bone marrow (BM) of contralateral and ipsilateral limbs (mean ± SEM, *n*=3 mice per group. Two-tailed unpaired *t* test). *$P$<0.05, **$P$ < 0.01, ***$P$ < 0.001.
